# Protocol for 3D Virtual Histology of Unstained Human Brain Tissue using Synchrotron Radiation Phase-Contrast Microtomography

**DOI:** 10.1101/2023.11.08.566183

**Authors:** Ju Young Lee, Sandro Donato, Andreas F. Mack, Ulrich Mattheus, Giuliana Tromba, Elena Longo, Lorenzo D’Amico, Sebastian Mueller, Thomas Shiozawa, Jonas Bause, Klaus Scheffler, Renata Longo, Gisela E. Hagberg

**Author notes:** Correspondence: **Ju Young. Lee**. Equal contributions as last authors (Renata Longo, Gisela E. Hagberg).

## Abstract

X-ray phase-contrast micro computed tomography using synchrotron radiation (SR PhC-µCT) offers unique 3D imaging capabilities for visualizing microstructure of the human brain. Its applicability for unstained soft tissue is an area of active research. Acquiring images from a tissue block without needing to section it into thin slices, as required in routine histology, allows for investigating the microstructure in its natural 3D space. This paper presents a detailed step-by-step guideline for imaging unstained human brain tissue at resolutions of a few micrometers with SR PhC-µCT implemented at SYRMEP, the hard X-ray imaging beamline of Elettra, the Italian synchrotron facility. We present examples of how blood vessels and neurons appear in the images acquired with isotropic 5 µm and 1 µm voxel sizes. Furthermore, the proposed protocol can be used to investigate important biological substrates such as neuromelanin or corpora amylacea. Their spatial distribution can be studied using specifically tailored segmentation tools that are validated by classical histology methods. In conclusion, SR PhC-µCT using the proposed protocols, including data acquisition and image processing, offers viable means of obtaining information about the anatomy of the human brain at the cellular level in 3D.

## Introduction

There is a great interest in studying the three-dimensional microstructure of human tissues, which is critical for improving our understanding of the spatial relationship between anatomical structures. Classical histology is the analysis of thin tissue slices that are stained and imaged using optical microscope. This is the gold standard for many biological fields, such as neuroscience. However, histology presents some disadvantages: during the sectioning deformations are introduced and it provides only 2D information. Advancement in image processing allows for correction of non-linear deformations to reconstruct 3D volumes (Adler et al., 2014, Amunts et al., 2020, Tendler et al., 2022). However, the histology slices are sometimes torn or folded, introducing discontinuities that may limit 3D reconstruction.

Ideally, tissue microstructure information could be obtained directly in 3D space. Light-sheet imaging is emerging as a sensitive and specific tool for volumetric imaging. However, the process of tissue clearing and labeling is required prior to imaging (Mai et al., 2023). Application of this technique can be challenging for large human samples (Park et al., 2022). While magnetic resonance imaging (MRI) and computed tomography (CT) are valuable sources for internal structure of the brain in clinical settings, they are limited in spatial resolution and image contrast.

In *ex vivo* MRI, where tissues are taken out of the body, the spatial resolution can be improved. In fact, *ex vivo* MRI of whole brain has been acquired with minimum voxel sizes of 100**^3^** – 400**^3^** µm**^3^**(Edlow et al., 2019, Shepherd et al., 2020). For smaller samples, smaller voxel can be achieved within feasible measurement time with high field scanners. For instance, 37**^3^** µm^3^ voxel size for samples with a diameter of 4 cm has been demonstrated (Tuzzi et al., 2020). Recent developments of dedicated hardware open up the prospect to perform MRI at the cellular level (Flint et al., 2020, Handwerker et al., 2020), albeit within a limited total spatial coverage of a few hundred micrometers. However, the magnetic properties of the tissue, which is reflected in MRI contrast, are influenced by the fixation procedure and the embedding media which limits reproducibility across tissue preparation pipelines (Birkl et al., 2016, Dusek et al., 2019, Nazemorroaya et al., 2022).

The image contrast in CT arises from the attenuation of X-ray beams as they pass through different materials. The attenuation depends on the material composition and the density. In soft tissue, there are no large differences between biological compartments, resulting in weak – or nearly absent-intrinsic contrast. Therefore, prior to CT imaging, the samples are usually injected with exogenous contrast agents that have densities higher than soft tissue. For example, in order to investigate the blood vessel structure, perfusion of contrast agents such as Microfil or Indian Ink into the blood vessels can be performed (Xue et al., 2014, Wälchli et al., 2021). Using conventional clinically available CT systems, typical voxel-sizes are 400**^3^** – 600**^3^** µm**^3^**, while the addition of photon-counting can push this value to 150**^3^** µm**^3^** (Wehrse et al., 2023).

Coherent X-rays can also yield contrast dependent on subtle phase shifts occurring in the tissue. Among such phase-sensitive techniques, propagation-based imaging (PBI), sometimes called free-space-propagation or single distance imaging, is available at several synchrotron light sources (Rigon et al., 2014). Phase shifts arise when X-rays with a certain degree of coherence pass through materials with varying refractive indices and can be interpreted as a local deformation of the X-ray wavefront. The phase difference between two X-rays passing at the interface between two compartments with different refractive indices, results in an interference pattern. This further develops as the beam propagates along an extended pathway free of obstacles.

In PBI, the sample-to-detector distance must be sufficient to allow the effects of subtle phase shifts caused by small structures to play out and be detected. The resulting image shows a sharp fringe edge enhancement at the tissue interfaces which is proportional to the Laplacian of the phase-shift (Peterzol et al., 2005). Through the application of a phase-retrieval algorithm, the fringes are compensated, and the original edge-enhanced image becomes an area contrast image. In case of small differences between the refractive indices, which is plausible in unstained tissue scanned in the near-field regime, the Paganin’s phase-retrieval algorithm can be employed (Paganin et al., 2002). Since the algorithm acts as a low pass filter, the final image has a higher signal-to-noise ratio than an attenuation image acquired without any free-space propagation would have. It is worth of notice that the phase-retrieval algorithm decreases the noise level of the image without affecting the spatial resolution (Brombal et al., 2020).

SR PhC-µCT images allow mapping the underlying anatomical structures, with high contrast-to-noise ratio despite the absence of large differences in electron density within the tissue. Hence, SR PhC-µCT is regarded as a suitable tool for virtual histology (Saccomano et al., 2018, Müller et al., 2021). Although it can be combined with injected contrast agents, these are not necessary per se to achieve high tissue contrasts. A wide palette of tissue preparations, like formalin-ethanol– or xylene-soaked tissue as well as paraffin embedded samples can be used for imaging with SR PhC-µCT (Rodgers et al., 2021).

Several studies have used SR PhC-µCT to investigate the microstructure of unstained human organs (Walsh et al., 2021, Einarsson et al., 2022, Lee et al., 2022). Some have acquired images from human brain samples with sub-cellular detail and successfully segmented neurons to study their three-dimensional spatial distributions (Hieber et al., 2006, Frost et al., 2023). SR PhC-µCT also has the potential for detecting pathology such as amyloid plaques and mineralized blood vessels (Astolfo et al, 2016, Toepperwien et al, 2020).

With experience of several beamtime sessions at the SYRMEP beamline of Elettra synchrotron, we have developed a pipeline that works reliably with unstained human brain tissue (Figure 1). We present a step-by-step guideline with emphasis on important decisions that need to be made at each step. We also describe how segmentation tools can be tailored to identify specific structures, and how these can be directly validated by classical histology methods applied to the same specimens. Segmentation methods were developed to identify blood vessels as well important biological substrates such as neuromelanin or corpora amylacea. Although the suggestions presented here are mainly relevant for the SYRMEP beamline, some of our proposed methods may be useful for SR PBI micro-CT performed at other synchrotron facilities. For example, we explored the transferability of some of our segmentation method to images obtained at other beamlines.

**Figure 1.**
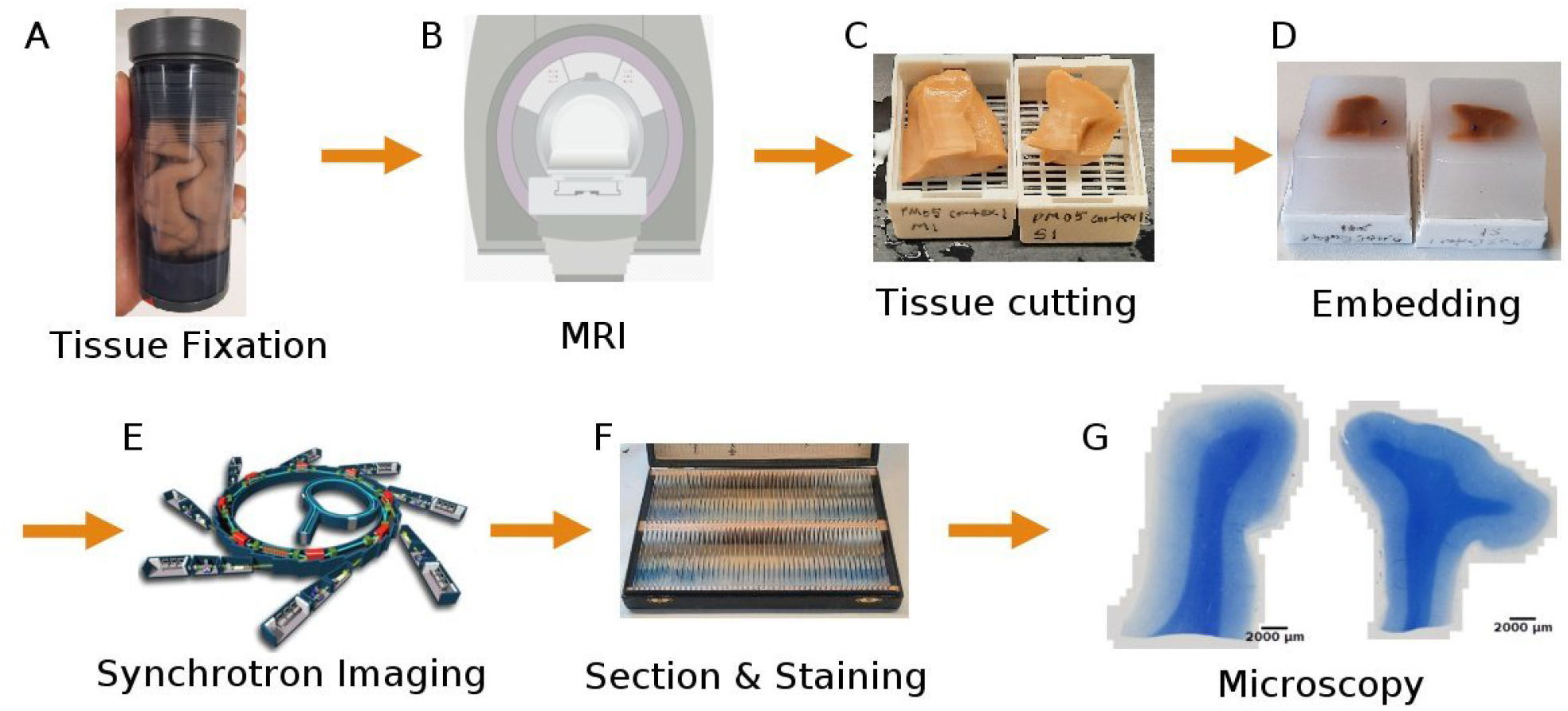
Complete workflow for the virtual histology of human brain tissue. (A) First, the target brain region is cut out and immersed in a fixative agent. The presented picture shows the cortical areas surrounding the central sulcus, which includes the primary motor and somatosensory regions. (B) If applicable, structural MRI may be acquired. (C) The tissue is cut into 1 cm thick sections to fit into the embedding cassettes. The left tissue is the hand area of the primary motor cortex and the right tissue is the hand area of the somatosensory cortex. (D) The tissues are embedded in paraffin and (E) brought to the biomedical imaging beamline of a synchrotron facility (F) After the beamtime, the paraffin blocks are sectioned into thin slices for (G) microscopy scanning. (Image credits for (E) is EPSIM 3D / JF Santarelli, Synchrotron Soleil)

## Protocol

### Step 1. Tissue preparation

The first step is to obtain *ex vivo* tissue. We obtained tissue from Tübingen University body donor program at the Institute of Clinical Anatomy and Cell Analysis. The body donors provided informed consent, in alignment with the Declaration of Helsinki’s guidelines for research, to donate their bodies for research purposes. The ethics commission at the Medical Department of the University of Tübingen approved the research procedure. It is also possible to obtain samples from brain banks that are already fixed and embedded (e.g. Netherlands brain bank).

### Step 1a. Dissection

Brain regions like the choroid plexus and pineal gland are likely to be calcified in aged subjects (Bukreeva et al., 2022, Junemann et al., 2023). Provided their anatomical locations are known, these areas should be cut out to minimize the risk of artifacts. For the X-ray energy spectrum optimal for soft tissue contrast, the presence of significant calcium content leads to pronounced streaking artifacts due to its higher absorption compared to the surrounding tissues (Orhan et al., 2020). An example of such effects caused by calcifications in the pineal gland is shown in Supplementary Figure 1.

### Step 1b. Fixation

After dissecting the targeted region, the tissues are either perfusion or immersion fixed using a fixative agent, for instance ethanol or perhaps more typically a formalin solution containing 4 % formaldehyde. Fixation time depends on the size of the sample. In case of human brain stems, we kept them in fixative solution for a minimum of 3 weeks, based on the experimentally determined diffusion-coefficient of the used fixative. If available, magnetic resonance imaging (MRI) can be scanned at this stage. In our case, we used a formalin solution that is optimized for *ev vivo* MRI at high magnetic field strengths (Nazemorroaya et al., 2022). Quantitative MRI can be used to check the advancement of the fixation process. Moreover, MRI is useful for calculating the shrinkage ratio introduced by paraffin embedding, which is the next step (Wehrl et al., 2015, De Guzman et al., 2016, Lee et al et al., 2022, Nazemorroaya et al., 2022)

The choice of the fixative agent will impact the tissue shrinkage ratio (Eckermann et al., 2021, Rodger et al., 2021) and tissue contrast in the resulting image (Strotton et al., 2018). For example, the fiber tract contrast is improved when tissue is fixed with 100 % ethanol compared to formalin (Rodger et al., 2021).

### Step 1c. Embedding

Different embedding materials may be used in accordance with the research scope (Strotton et al., 2018). We used formalin-fixed paraffin-embedding (FFPE) which is suitable for histology afterwards. Many researchers have used cylindrically shaped biopsy punches with diameter and height of a few millimeters (Hieber et al., 2016, Töpperwien et al., 2020, Saiga et al., 2021). In contrast, we used larger samples than usual size for routine histology, with width and height of 2 ∼ 3 cm. Therefore, extra care was taken during the paraffin embedding process. The automated embedding station normally employed for smaller samples may lead to incomplete paraffin penetration and large deformation such as dented surfaces (Supplementary Figure 2) (Zhanmu et al., 2020).

In order to reduce SR PhC-µCT image artifacts, we advise to prevent air bubbles from being trapped in the paraffin blocks. Air-tissue boundaries create strong edge-enhancements that will turn into strong image artifacts due to large difference in their refractive indices (Supplementary Figure 1). In order to minimize the occurrence of entrapped air bubbles, we placed the sample in vacuum during the paraffin embedding process. The use of vacuum pumping during the embedding process is a routine practice in resin embedding for electron microscopy and is therefore known not to damage the tissue. Some residual air bubbles may remain even with repeated degassing processes (Brunet et al., 2023). Other approaches such as keeping the paraffin wax at 60 degrees for a long time could be considered to further improve this process (Zhanmu et al.,2022).

### Step 2. Imaging at the synchrotron facility

Elettra is a third-generation synchrotron located in Trieste, Italy. It is equipped with 263 m long electron storage ring that supplies several beamlines, including SYRMEP (SYnchrotron Radiation for MEdical Physics) beamline. The beamline bending magnet provides polychromatic light in the energy range from ∼8.5 to 40 keV. The X-ray source is located at > 20 m from the experimental station, thus the impinging beam has high spatial coherency, allowing for propagation-based phase-contrast imaging. In our experiment, the polychromatic (white) beam mode, rather than monochromatic beam mode, was used to maximize the photon flux. With a silicon filter, the average energy of the beam was 20 keV (Dullin et al., 2021).

Inside the experimental station of the SYRMEP beamline, the sample is placed on a rotating stage at a 23 m distance from the light source (Figure 2). The beam height at this position is approximately 4 mm. Even if a larger footprint of the beam can be used for image acquisition, our recommendation is to limit the beam size to the Full-Width-Half-Maximum. It will allow to obtain a more homogeneous signal-to-noise ratio within images of the same dataset. Acquisitions were performed with a 2048 x 2048 sCMOS detector with a physical pixel size of 6.5 μm. The used detector was coupled with a high-numerical aperture optic allowing to select effective pixel sizes between 0.9 um and 5 um. The X-ray was converted into visible light through a gadolinium gallium garnet Eu-doped (GGG:Eu) scintillator screen (Brombal et al., 2020, Donato et al., 2022). The pixel size determines the vertical field-of-view (FOV). If the height of the sample is bigger than the beam height as shown in Figure 2, the entire vertical length cane be covered by multiple tomographic acquisitions (further elaborated in Step 2d). In practice, we opted for two beamline setups. One encompassing larger pixels of ∼ 5 µm x 5 µm to increase the FOV of each acquired image, another with smaller pixels of ∼ 1 µm x 1 µm to allow observations of finer details (Figure 3, Table 1). Since the phase-shift pattern depends on the sample-to-detector distance, this parameter has to be optimized according to the detector pixel size. Below, we explain additional factors to consider during the beamtime in detail.

**Figure 2.**
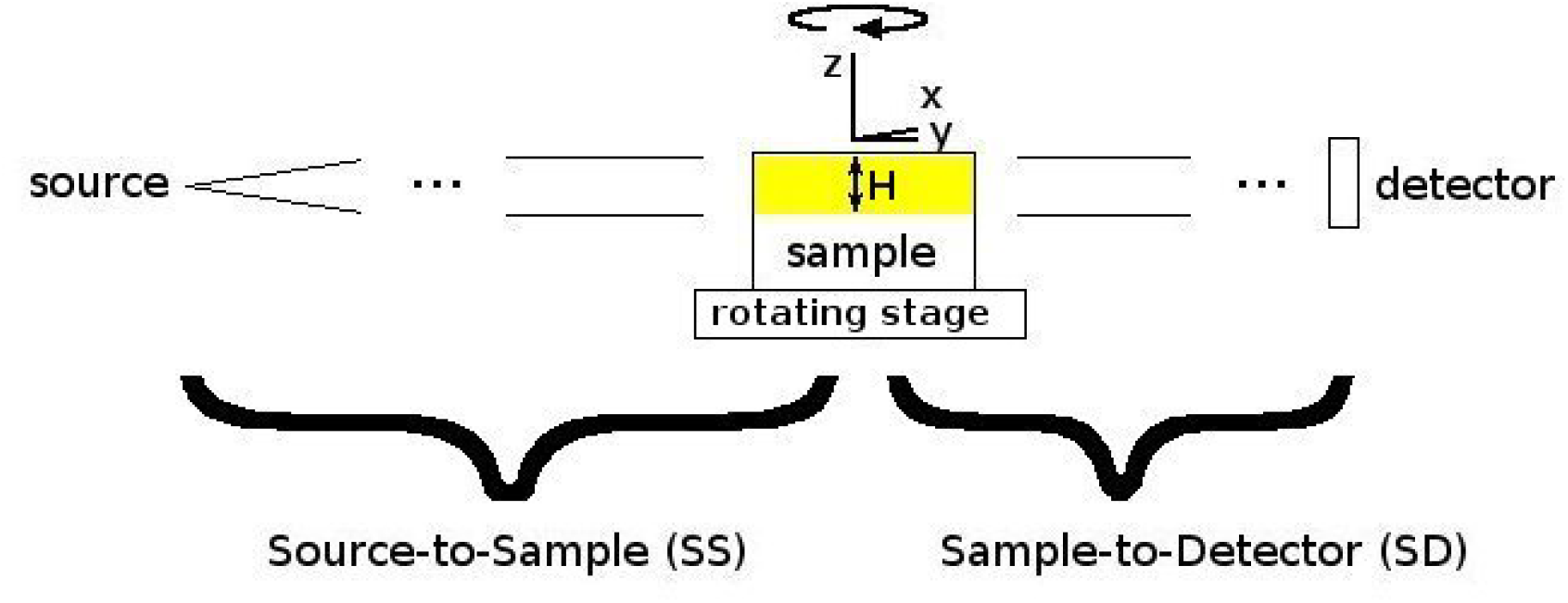
Schematic picture of the beamline setup. To obtain tomographic reconstructions, the sample is rotated around the vertical Z axis. H is the source height. scintillator screen (Brombal et al., 2020, Donato et al., 2022). The pixel size determines the vertical field-of-view (FOV). If the height of the sample is bigger than the beam height as shown in Figure 2, the entire vertical length cane be covered by multiple tomographic acquisitions (further elaborated in Step 2c).

**Figure 3.**
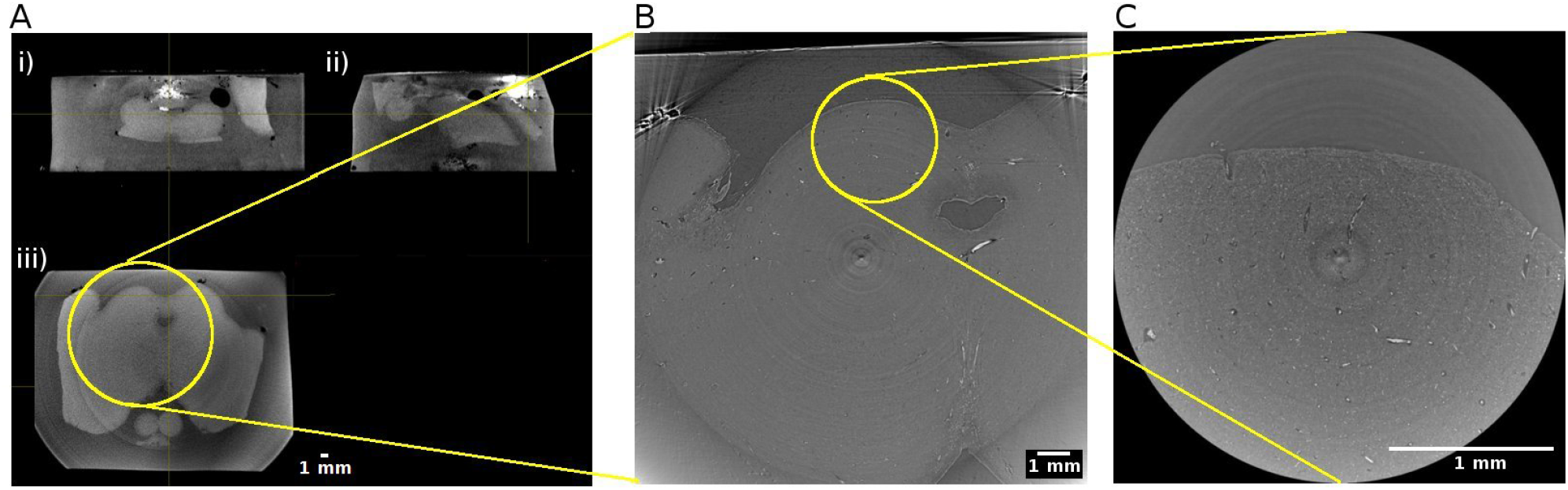
Hierarchical imaging of the midbrain of the human brainstem. (A) Cone beam imaging from Tomolab (Elettra, Trieste) covers the entire volume of the sample and is acquired with a voxel size of 20 µm. Orthogonal views showing i) coronal; ii) sagittal; and iii) axial views through the sample. Axial views from synchrotron radiation phase-contrast microtomography acquired with 5 µm isotropic voxel (B) and 1 µm isotropic voxel (C). Yellow circles indicate the matching regions when going from the lower to the higher resolution measurements.

**Table 1.** Hierarchical imaging beamline setup for propagation-based phase-contrast microtomography. The specifications apply to experiments at SYRMEP beamline at Elettra synchrotron facility. The details may vary between sites.

|  | Sample-to-detector distance | Reconstruction window | Acquisition time |
| --- | --- | --- | --- |
| (5 $\mu\text{m}$)<sup>3</sup> voxel | 900 mm | Lateral: $\sim 17$ mm<br>Vertical: $\sim 4$ mm | 20 min |
| (1 $\mu\text{m}$)<sup>3</sup> voxel | 200 mm | Lateral: $\sim 3$ mm<br>Vertical: $\sim 2$ mm | 30 min |

### Step 2a. Pixel size and sample-to-detector distance selection

We utilized the PBI, which is applicable in the near-field region. This condition is met for large Fresnel numbers *N_F_ = d_eff_^2^/(λR_eff_) >> 1*, where *d_eff_*is the effective pixel size, *R_eff_* is the sample-to-detector distance multiplied with the square of the magnification factor (the ratio of the source to detector over the source to sample distance) and *λ* is the X-ray wavelength. In experimental practice, this condition is often relaxed to allow N_F_ to be closer to 1. Once the detector pixel size is determined, this condition places a constraint on the maximum propagation distance. In the context of FFPE human tissues, the signal-to-noise ratio in reconstructed images is optimized for *N_F_* > 2 (Donato et al., 2022).

In our measurements, a sample-to-detector distance of 90 cm was used for the ∼ 5 µm pixel size and 20 cm was used for the ∼ 1µm pixel size. In future studies, these distances can be optimized in order to obtain better signal-to-noise ratio (Donato et al., 2021)

When imaging a large sample like parts of the human brain, it is strongly recommended to acquire data using at least two pixel size settings. Images with a larger pixel size will have a larger field of view, covering several landmarks of the sample. Images with a smaller pixel size will be a zoomed in image with a smaller field of view. This method is sometimes referred to as hierarchical imaging or multi-scale imaging (Walsh et al., 2021). Even if the research question only concerns the microstructure of a particular part of the tissue, it is always advisable to acquire images with a larger field of view as a reference. Otherwise, it may be challenging to infer the exact location of the high spatial resolution image and to position the acquired information within its exact anatomical context (Figure 3).

### Step 2b. Selection of the scintillator

High resolution detectors for soft tissue CT are usually indirect conversion type, requiring incident X-ray beam to be converted into visible light. Typically, a scintillator screen is used for converting the beam. Synchrotron beamlines commonly employ optical systems that allow for selecting different scintillator types (Lecoq et al., 2016). When selecting a scintillator, users need to consider the conversion efficiency of the X-rays to visible light and its impact on spatial resolution. Thicker scintillators exhibit higher efficiency, but may also increase signal blurring, consequently reducing spatial resolution. On the other hand, thinner scintillation materials better preserve spatial resolution but have lower efficiency, requiring longer exposure times. Balancing between acquisition time and the desired resolution is essential. At SYRMEP beamline, we used GGG:Eu scintillators with thickness of 45 µm for a voxel size of 5^3^ µm^3^ and 17 µm thickness for a voxel size of 1^3^ µm^3^ (Supplementary Figure 3).

### Step 2c. Planning for sample coverage

Based on the decided pixel size, it is possible to estimate the size of the FOV. Typically, the considered reconstruction window will have sizes determined by the width of the photon detector. In order to enlarge the horizontal FOV, the rotation axis can be placed off-center, i.e. at one edge of the camera FOV. This approach is known as extended FOV (or half-acquisition) method (Wang et al., 2002). This method requires a dedicated pre-processing, called stitching of the sinogram prior to reconstruction which is explained in Step 3a.

If the extended FOV is smaller than the brain sample under investigation, it is helpful to plan ahead how to position the FOV of each acquisition. We cut out circular shapes from an adhesive tape with diameter corresponding to the size of the FOV. For the acquisitions with voxel sizes of 5**^3^** µm**^3^**, the diameter was about 1.7 cm. These circles were placed on top of the specimen to mark each FOV. The number of circles required then gave information about how many scans were needed to obtain full coverage the region of interest, also known as mosaic CT. As a reference to position the sample, a marker is placed above the center of each FOV for ‘Step 2f’ (Figure 4).

**Figure 4.**
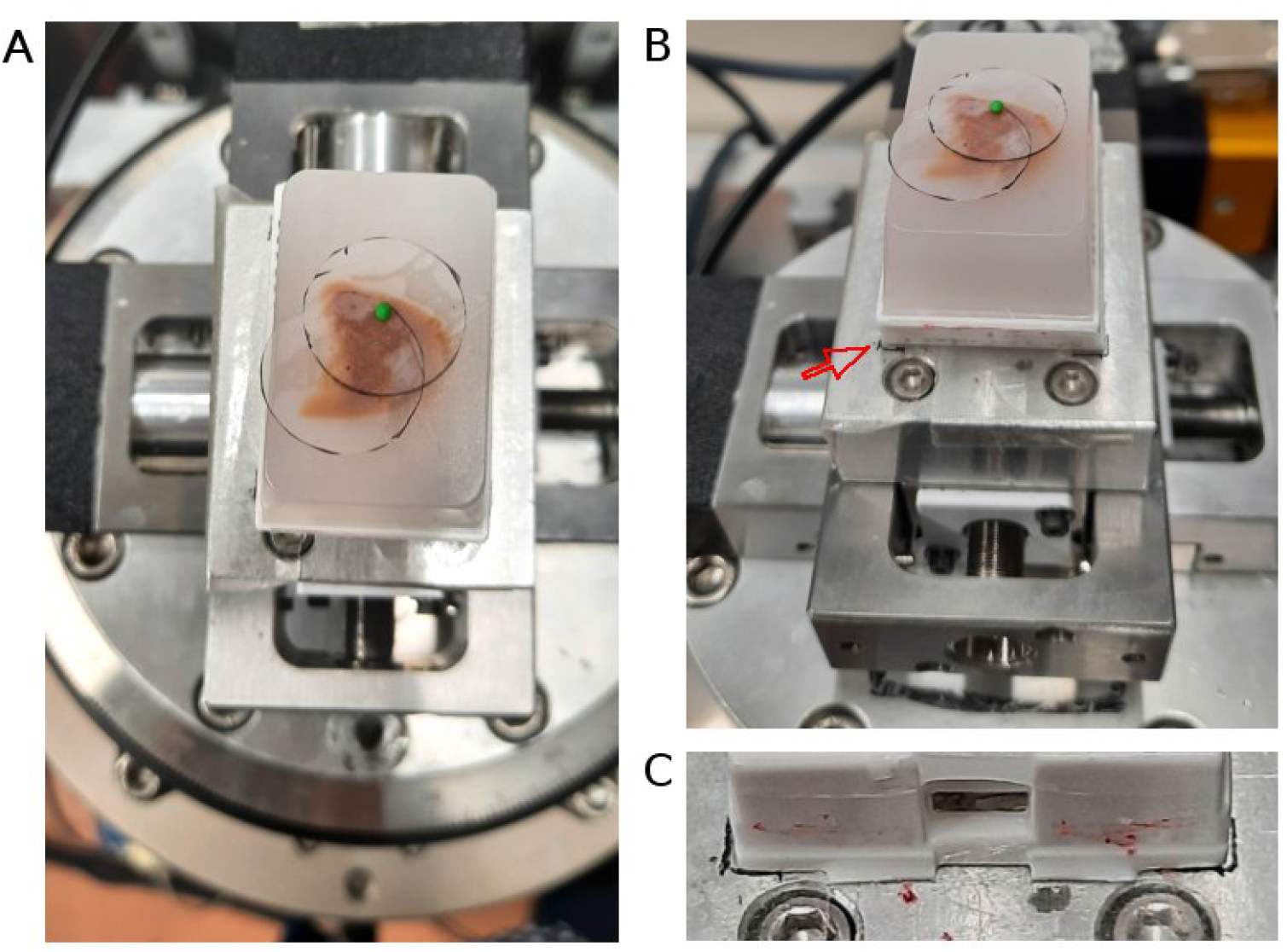
Sample positioning and coverage planning at SYRMEP beamline experimental station. (A) A paraffin embedded brain sample was placed on the rotator. The coverage of the measurement was estimated by a transparent plastic film placed on the sample surface. A small green polymer clay, which serves as a radiopaque marker, was placed in the center. (B) The same setting as panel A with picture taken from slightly lower angle. The red arrow points to the marked position drawn on the adhesive tape. (C) A close up image showing the boundary of the plastic cassette drawn as a black line.

For our FFPE samples, the vertical sample length was around 1 cm. Thus, multiple vertical acquisitions were needed for each specimen. Working with 5 µm pixel, the vertical field of view was limited to 4 mm, corresponding to the available beam height. For 1 µm pixel setting, the vertical FOV of the detector was ∼ 2 mm,. In order to collect data from the whole cylindrical volume planned in ‘Step 2c’, several scans should be made at different vertical positions of the sample. The volumes should be partially overlapping to enable stitching process in Step 3e. For example, sample shown in Figure 4 required two vertical steps for both lateral FOV, amounting to 4 scans in total. At SYRMEP beamline, vertical steps can be automatized using a dedicated script that control the sample position.

### Step 2d. Sample placement on the rotator stage

We used double sided adhesive tape to stabilize the sample and mark the position on the sample holder that is placed on top of the rotator. It is useful to mark the position of the sample on the sample holder, so that the samples can be placed in roughly the same position for measurements with different sample-to-detector distances facilitating their 3D registration (Fig. 4).

### Step 2e. Acquisition

A radiopaque marker, such as polymer clay, can be played above the sample facilitating the precise alignment of the area to visualize within the camera FOV. At the SYRMEP beamline the sample centering is typically performed by using two orthogonal precision stages (micrometer linear motors) placed above the rotator. It is highly recommended to remove the markers before launching the acquisition, as they can be the source of strong streak artifacts that can impair the visibility of several slices on the sample surface. Otherwise, users should make sure that the FOV doesn’t include the markers to avoid artifacts.

Each acquisition is composed of dark image (an image acquired without x-ray beam to measure the noise of the detector), flat image (an image acquisition without the sample between beam and detector), and projection measurement. We acquired 3600 projections across 0-360 degree in half-acquisition modality. The exposure time ranged between 150 – 200 ms.

Prior to launching the measurements, it is crucial to check for any damage or scratches on the scintillator that may result in abnormal pixel signal. Over the course of long measurement, scintillators may accumulate dust, leading to the generation of highly intense pixels in the projection that are difficult to normalize during the flat-field correction (Step 3a). These pixels can subsequently be the source of ring artifact. To mitigate these issues, users should check the flat field regularly (Figure 5).

**Figure 5.**
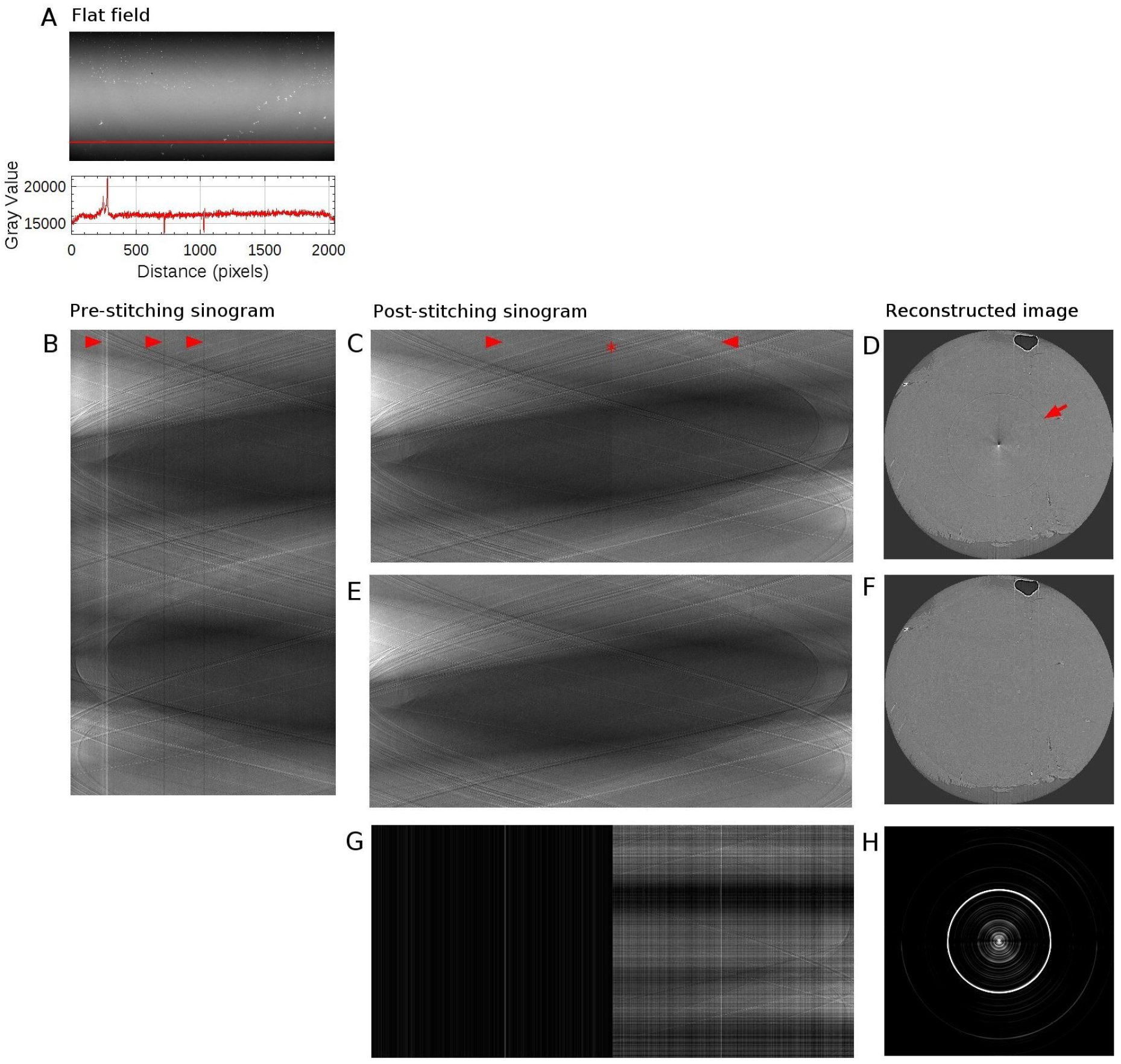
Sinogram stitching and ringing artifact removal. (A) The above image is a flat field image (matrix size: 1000 x 2048). The detector matrix size is 2048 x 2048, but the edge rows are excluded in order to use beam within its full-width-at-half-maximum only. The bottom plot shows the intensity profile along the red vertical line. (B) Sinogram with a sample in place, at the level of the red vertical line, measured with half-acquisition mode (3600 x 2048). 3600 projections were acquired over 360° with offset center-of-rotation. Red arrows point to stripe artifacts which coincides with the intensity variation in the flat field image (C) Stitched sinogram (1800 x 3732). Projections obtained at rotation angles 1-180° have been stitched together with angles 181-360° at the point indicated by a red star to yield the equivalent 180° sinogram. Stripe artifacts, occurring in agreement with variation in the flat field image, have been indicated by red arrows. (D) Reconstructed image from the sinogram shown in panel C. Ringing artifact is indicated with a red arrow. Artifacts at the center of the image is due to half-acquisition mode. (E) Stitched sinogram (1800 x 3732) obtained with ring removal filter and line-by-line normalization. (F) Reconstructed image from the sinogram shown in panel E. (G) Intensity difference between panel C and E. (H) Intensity difference between panel D and F. The brightness and contrast were adjusted to better visualize the difference in panel G and H. holder, so that the samples can be placed in roughly the same position for measurements with different sample-to-detector distances facilitating their 3D registration (Fig. 4).

### Step 3. Reconstruction

A standardized reconstruction pipeline for propagation-based phase-contrast CT includes a pre-processing step for image normalization also known as flat-field correction, phase retrieval, potential filtering for ring removal and a reconstruction algorithm such as filtered back projection. In this section, we demonstrate how each process of the reconstruction steps can influence the final result.

The 3D reconstruction procedure (Step 3a – Step 3c) was done with SYRMEP Tomo Project (STP) software suite (Brun et al., 2015, Brun et al., 2017), which is developed based on the Astra toolbox (Van Aarle et al., 2015). At our home institution, we used a workstation with 64 GB memory, 12 GB of graphic memory, and 12 physical CPU cores. A raw projections datasets of size around 10 GB required 1 hour from preprocessing to reconstruction with STP software (Table 2). The reconstruction process from flat-fielding correction to image reconstruction can be automatized once the parameters for each step are defined. However, the computation time depends on the hardware resources (CPU and GPU).

**Table 2.** Computation time. We used a computer with 64 GB memory with 12 physical CPU cores.

|  | Pre-process | Phase retrieval | Reconstruction |
| --- | --- | --- | --- |
| (5 µm)<sup>3</sup> voxel image | 10 min | 30 min | 25 min |
| (1 µm)<sup>3</sup> voxel image | 1 hr | 1 hr | 2 hr |

Raw data from each acquisition can consist of tenths of GB. They are archived at Elettra server for several years and can be retrieved if requested. The output of the STP software is a stack of reconstructed slices in 32-bit TIFF file format. 1 µm isotropic voxel images scanned with half acquisition mode can amount to over 100 GB after reconstruction. Typically, in 3 ∼ 5 days long beamtime, several TB of data can be produced.

For post-processing steps (Step 3d – Step 3f), we used a widely used software in the biology community, called ImageJ (Schneider et al., 2012, Schindelin et al., 2012).

### Step 3a. Sinogram

If the half acquisition method was used, the overlap should be estimated to perform sinogram stitching. The overlap can be estimated either using a Fourier-based algorithm (Vo et al., 2014) or by visual assessment. Typically, we first performed stitching using the algorithm and visually inspected the resulting sinogram. If the stitching result needed additional adjustment, we changed the parameter in small steps until the result improved. Usually, the overlap needed small correction for each vertical steps of the same lateral FOV.

Imperfect scintillators or defective pixels can be the source of ring artifacts in the reconstructed images. There are several ring artifact removal filters that can be tuned and tested. The strength of the ring removal filter can significantly reduce the artifacts, but can introduce blurring in the image. An alternative approach is to acquire images by introducing random shifts of a few pixels of either the detector or the sample during acquisitions, transforming ring artifact into dispersed noise (Liu et al., 2023). This approach requires projections in the step-and-shoot mode, thus increasing the acquisition time.

When the FOV is significantly smaller than the entire sample, it is recommended to use padding on the sinogram in order to reduce the local tomography artefact (Marone et al., 2010).

To compensate for imperfect flat-fielding due to factors such as beam instabilities, or time varying detector properties, users may consider using dynamic flat-field correction (Van Nieuvenhove et al. 2015).

### Step 3b. Phase retrieval

In propagation based SR PhC-µCT, the phase signal is retrieved from the acquired projections prior to the 3D reconstruction. A widely used phase retrieval algorithm is the one developed by Paganin and colleagues (2002). The algorithm assumes that the sample under investigation is homogeneous and that the data is acquired with monochromatic radiation. It uses the ratio between the real (δ) and the imaginary part (β) of the refractive index as a parameter. For non-homogeneous objects and polychromatic radiation, a common experimental practice is to use the algorithm parameters δ/β as a tuning parameter to regulate the strength of the filter and thus the blurring introduced in the image (Strotton et al., 2018). In practice, the algorithm acts like a low pass filter in the spatial frequency space. For FFPE brain samples, we used the δ/β in the range between 20 and 50.

### Step 3c. 3D Reconstruction

In this step, the matrix of the projections is used to calculate tomographic images. The STP software suite provides several reconstruction filters. This choice depends on the specific imaging application, as well as the desired trade-off between image sharpness and noise reduction. For this application, filtered back projection algorithm was applied using Shepp-Logan filter (Brun et al., 2017). This algorithm is known to be fast and efficient when applied to datasets with good signal-to-noise ratio and a large number of projections. A circularly shaped reconstruction window can be applied to limit the result within the FOV (as shown in Fig. 3C)

### Step 3d. Background removal

Due to the so called ‘beam hardening’ effect, images can exhibit a cupping artifact, which yields brighter intensity in the periphery than the inner region. There are several techniques to remove such artifacts. For example, we implemented slice-by-slice second degree polynomial fitting with the Xlib plugin (Münch, 2015) (Figure 6).

**Figure 6.**
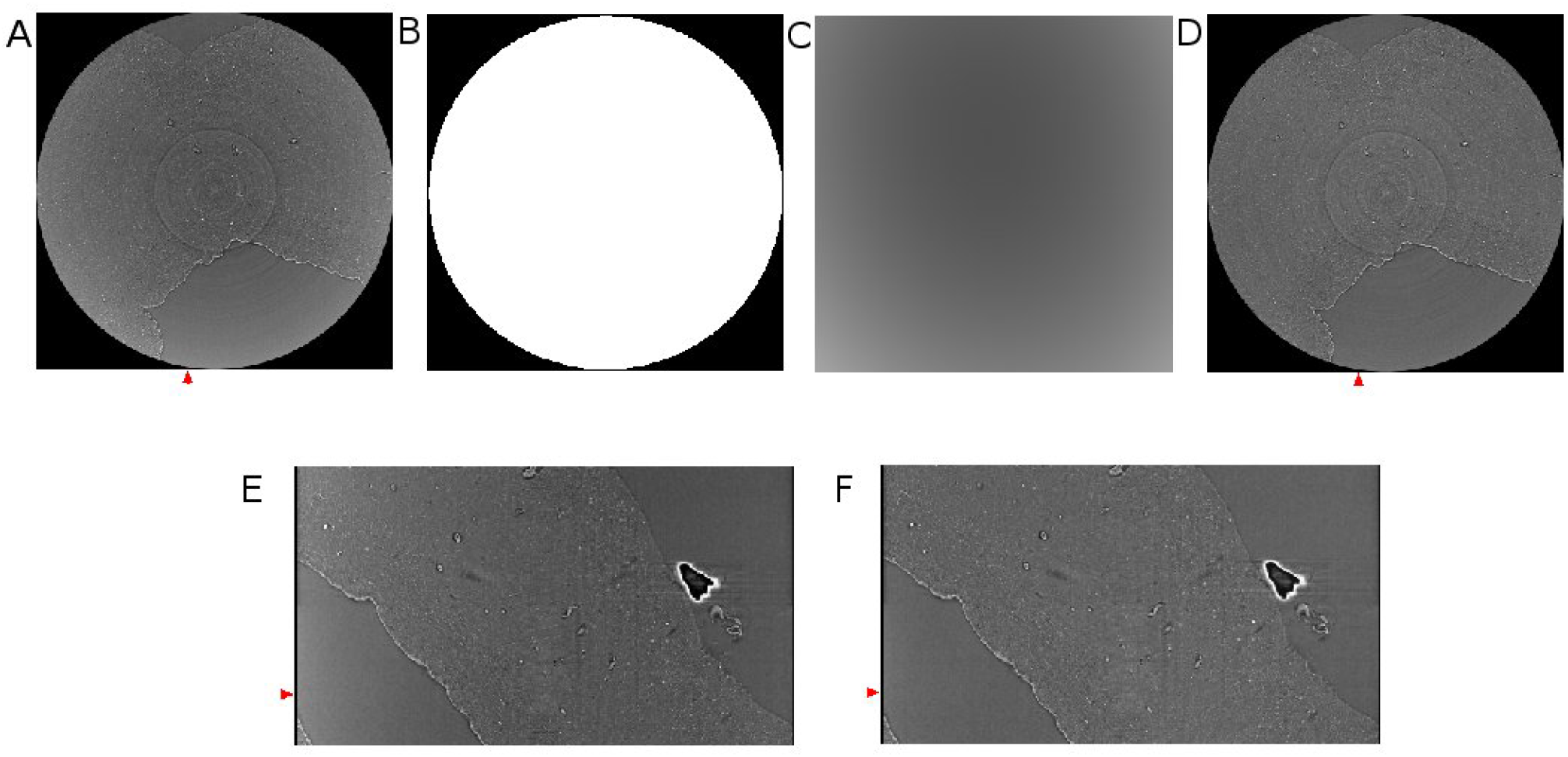
Background removal using polynomial fitting. (A) Reconstructed horizontal image showing cupping artifact. (B) Mask for background removal. (C) Background estimated using second degree polynomial fitting. (D) Corresponding slide shown in panel A after background removal. (E) A vertical slice of the image without background removal, taken from the region annotated with a red arrow in panel A. (F) A vertical slice of the image after background removal, taken from the region annotated with a red arrow in panel F. Red arrows in (E) and (F) match the position of the slides shown in panel (A) and (D).

### Step 3e. Stitching 3D reconstructed image

This is an optional step for users who want to combine neighboring volumes into a single volume. The pairwise stitching function (Preibisch et al., 2009) is a useful tool for merging 3D images. This function uses the Fourier transform to calculate the translation matrix from the frequency domain. For the function to perform well, there should be enough overlap between two images that need to be stitched together.

This algorithm can also be used for registering the high resolution image to the low resolution image. For example, we used this function to register the 1 µm isotropic voxel size images to the 5 µm isotropic voxel size image. The higher resolution images should first be downscaled to match the voxel size of the lower resolution image.

### Step 3f. Image Segmentation

In order to investigate the biological structures systematically and perform any volumetric quantitative measure, it is desirable to segment them. Manual segmentation requires a trained eye and a lot of time (Saiga et al., 2021). We suggest two segmentation approaches to automatically identify biological structures in SR PhC-µCT images: an intensity-based approach and an edge-based approach. These pipelines can be implemented with ImageJ plugins specialized for 3D image processing such as 3D ImageJ Suite (Ollion et al., 2013) and MorphlibJ (Legland et al., 2016). An example of a segmentation result using these methods are presented in Figure 7. These methods do not require training data or as much computational power as machine learning based approaches (mentioned in discussions).

**Figure 7.**
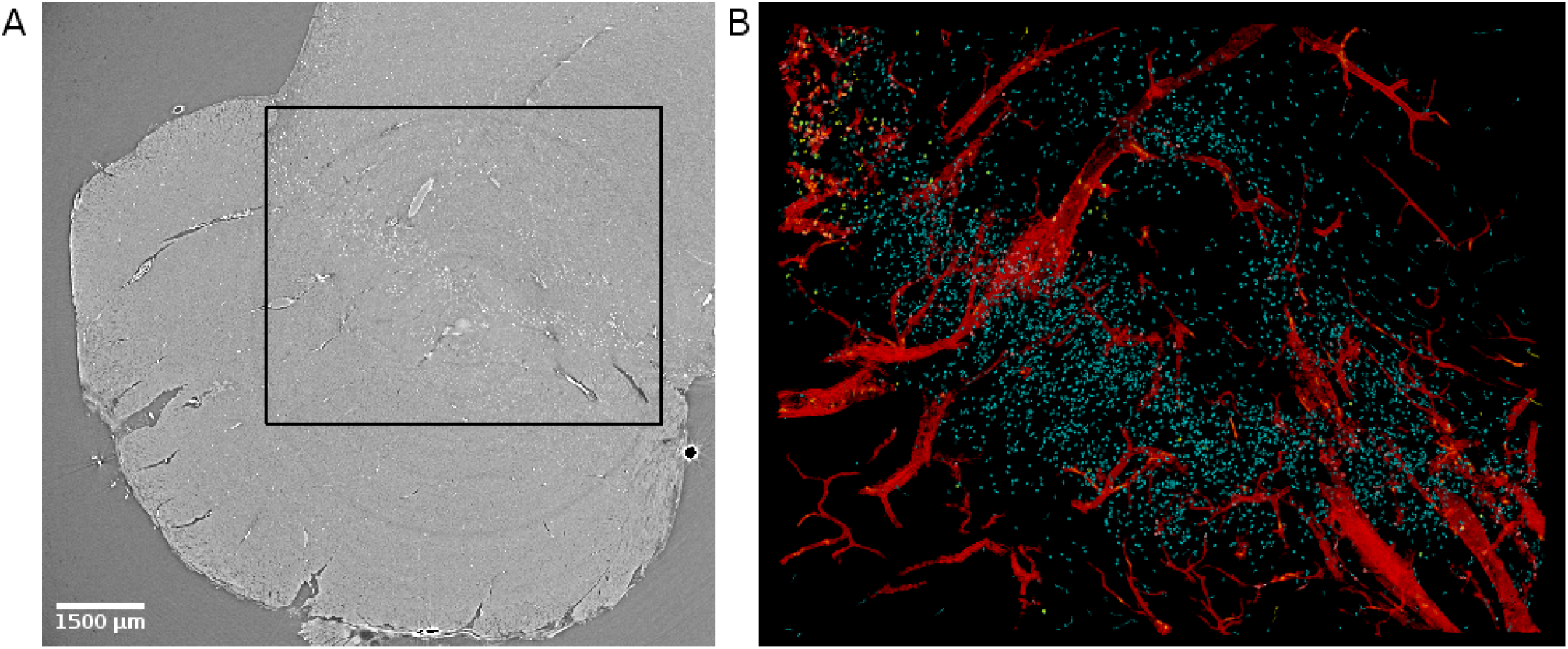
Virtual histology of the substantia nigra of the human brain stem. (A) Synchrotron radiation phase-contrast microtomography obtained with 4.74 µm isotropic voxel size. (B) Segmented results from the region indicated with a black rectangle in panel A. Blood vessels (in red) were segmented using an edge-based approach (Lee et al., 2022). Corpora amylacea (yellow) were segmented using an intensity-based approach (Lee et al., 2023) and a similar approach was used to identify neuromelanin (cyan). Corpora amylacea are clustered in the top left corner.

The intensity-based approach relies on the attenuation of X-ray dependent on the density and composition of the materials. Therefore, it is useful for identifying structures containing elements with high atomic weight (e.g. neuromelanin with iron) or dense structures (e.g. corpora amylacea). After applying a threshold to the image, the resulting image will be a binary image with many connected components. Morphological features of the connected components such as size and sphericity can be used to increase the accuracy of the segmentation (Lee et al., 2023).

The edge-based approach is suitable for detection of large connected structures such as blood vessels. Typically, a denoising process such as median filter precedes the edge detection algorithm. A median filter can remove salt-and-pepper noise while preserving edge structures. Deriche-Canny edge detection followed by a flood-filling process was proposed in our previous work (Lee et al., 2022).

With high resolution images amounting to large data sizes, users should consider working with down-sampled images. If the structure of interest is large enough, down-sampling can be used to reduce the computation time.

### Step 4. Validation with histology

Histology is essential for validating the observed structure in SR PhC-µCT as known biological entities. The advantage is that histology can be performed on the same FFPE samples as those used for SR PhC-µCT. Using a rotational microtome (Leica, Wetzlar, Germany), we cut the paraffin block into ∼ 10 µm thick sections. These sections can be stained for investigating specific research topics.

It has been shown that X-ray exposure during SR PhC-µCT does not hinder subsequent histological analysis (Saccomano et al., 2018). We have tested four types of classical staining methods that reliably work, including hematoxylin eosin stain and luxol fast blue stain. Table 3 summarizes how color appearance can be interpreted. In our experiments, immunohistochemical staining was not reliable, presumably due to prolonged formalin fixation and depending on the epitope. Among the antibodies that we have tested, anti-CD34 (a marker for endothelial cells) and anti-Aquaporin 4 antibodies were successful but ERG (ETS-Related Gene) antibodies were not. In typical research settings, small pieces of tissue is fixed in formalin for about 24 hours. Since the samples we used were of ∼2 cm^3^ volume, longer storage in fixative solution was necessary. Commercial antibodies are likely not optimized for tissues stored in fixative solution for longer times. Furthermore, the differences in perfusion versus immersion fixation is known to influence antibody binding (Woelfle et al., 2023)

**Table 3.** List of histology methods. These techniques successfully worked for brain tissue that were scanned with synchrotron radiation imaging. The right column describes how the biological structures appear with the corresponding staining.

| Method | Appearance |
| --- | --- |
| Hematoxylin – Eosin | Cell nuclei: blue<br>Cell body: violet |
| Luxol Fast Blue – Cresyl Violet | Myelin: blue<br>Cell body: violet |
| Periodic Acid Schiff -Hematoxylin | Polysaccharide: Violet<br>Cell nuclei: blue |
| Elastic van Gieson | Collagen: brown |

3D reconstruction of histological slides by serial alignment is very challenging due to non-linear deformation. Several methods have been proposed (Amunts et al., 2013, Haddad et al., 2021, Howard et al., 2023). Besides, the registration of microscopy images to µCT images is also not trivial, since microscopy images are not isotropic as in microtomography (Albers et al., 2021). We recommend using histology for characterizing biological components. How to do so is explained below.

### Examples of features identified from unstained human brain

In this section, we present biological features that can be extracted from SR PhC-µCT acquired from unstained human brain samples. Figure 7 demonstrates how multiple features can be extracted from a single PhC-µCT image. An automated segmentation pipeline was used to extract blood vessels, neuromelanin and corpora amylacea as explained in ‘Step 3f’. Each feature will be further discussed in the following paragraphs.

We mainly present our own measurements from FFPE samples acquired at SYRMEP beamline. For comparison, images from other open source data acquired at different synchrotron facilities are shown when applicable (Walsh et al., 2021, Eckermann et al., 2021: PNAS). Note that variations in tissue preparation such as hydrated/dehydrated, embedding material, frozen, can change the contrast (Töpperwien et al., 2019, Eckermann et al., 2021, Rodger et al., 2021).

As we are not introducing an external contrast agent, the SR PhC-µCT signal relies on the intrinsic density and tissue composition (Piai et al., 2019). Biological structures that are dense or contain materials with high electron density will lead to increased attenuation. Increased attenuation in a region implies that the structure is 1) composed of elements with high atomic weight (e.g. iron, zinc) and/or 2) are very densely packed.

### Blood vessels

It is challenging to investigate the blood vessel structure using traditional 2D histology, where the tissues are cut into thin slices before imaging. SR PhC-µCT can provide 3D isotropic images that preserve the continuity of the blood vessel. When exploring SR PhC-µCT data by simple visual inspection, the trajectories of large blood vessels are prevalent features one can easily notice (see Video 1).

Figure 8 shows examples of different brain regions measured with SR PhC-µCT, paired with segmented blood vessel structures. The brain stem acquired with 5 µm isotropic voxels (Fig. 8A) was processed using the edge-based vessel segmentation method detailed in Lee et al., (2022). The resulting vessel map (Fig. 8B) matches the known anatomy of the blood vessel in the region as previously reported (Naidich et al., 2009). For whole brain data, it is possible to follow the trajectory of the oxygenated blood from the heart by leveraging on the known blood source to the brain, called circle of Willis (Fig. 8C). We used a semi-automated seed-based segmentation method with the ITK-SNAP software (Yushkevich et al., 2006) to trace the arteries of a hemisphere by following arteries branching off from the circle of Willis (Fig. 8D). Lastly, SR PhC-µCT of the occipital cortex (Fig. 8E) was processed with an edge-based segmentation method to reveal the penetrating vessels running orthogonally to the surface of the cortex (Fig. 8F).

**Figure 8.**
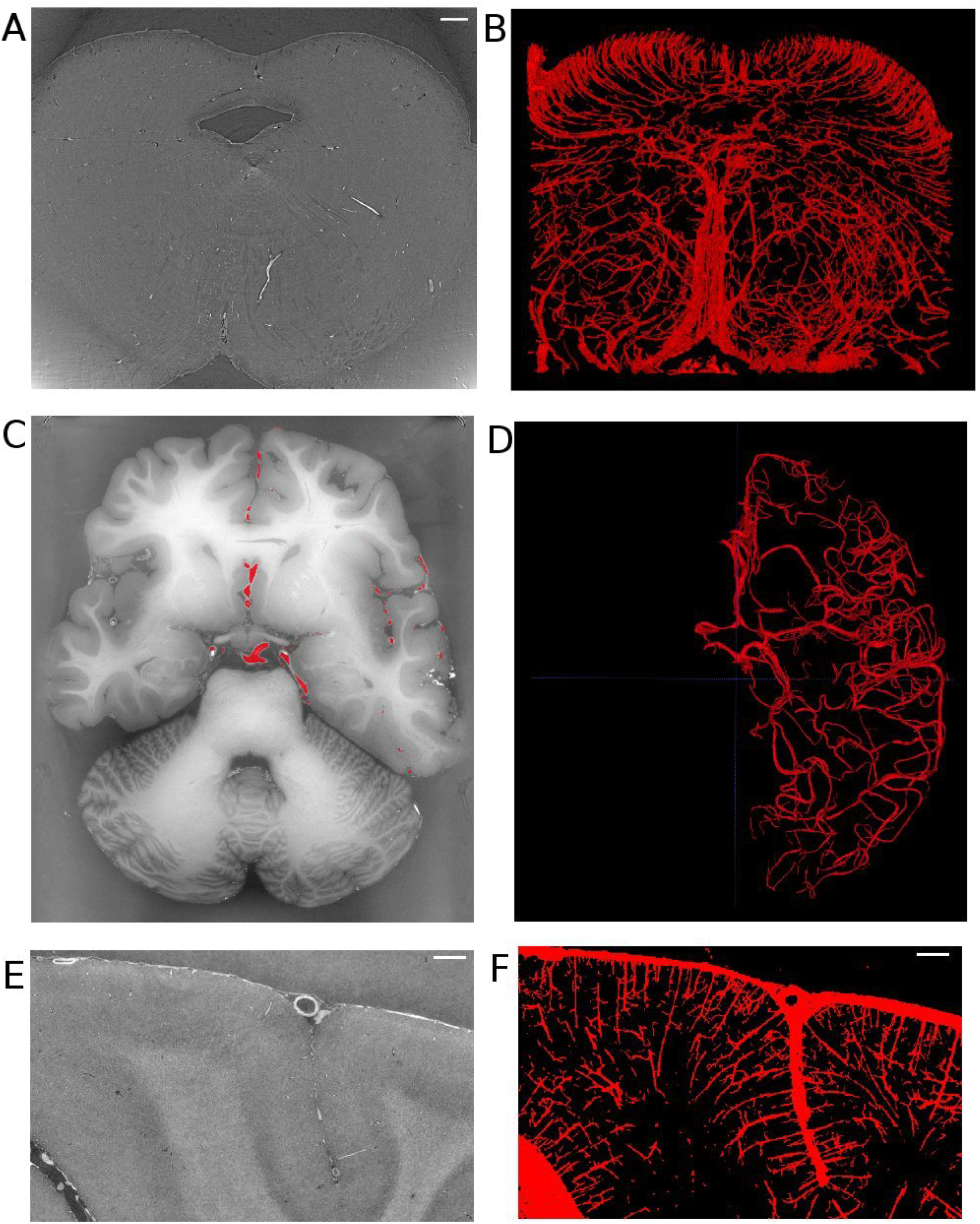
Vessel extraction from synchrotron radiation phase-contrast microtomography. (A). Axial view of the human brainstem. (B). Vessel segmentation using edge-based automated segmentation from the region shown in A (C). Axial view of the whole brain. (D) Result of artery tracing of one hemisphere using seed based semi-automated segmentation with ITK-SNAP. (E) Coronal view of the occipital lobe. (F) Vessel segmentation result using edge-based segmentation from the region shown in panel E. Scale bars in A, E, F are 1 mm in length. (C-F) are based on open source human organ atlas data (Walsh et al., 2021). Panels B and D are 3D rendered views of a 1.5 mm thick slab and a hemisphere, respectively. Panel F is a maximum intensity projection across 1 mm depth. DOI identifier for (C) is 10.15151/ESRF-DC-572252655 and (E) is 10.15151/ESRF-DC-572253460.

Note that the appearance of blood vessels may be different in hydrated tissues (Ekermann et al., 2021: Biomed.Optics).

## Neurons

Some large neurons could be visualized with SR PhC-µCT. Figure 9 shows the large neurons of the mesenphalic trigeminal nucleus acquired with 5 µm and 1 µm isotropic voxels. Individual cells are sharply delineated only in the 1 µm isotropic voxel image. Figure 10 shows examples of smaller neurons observed with 1 µm isotropic voxel size. Histology of the matching region was used to validate these structures as neurons (Fig. 10A-D). About the center of the cell body, the nucleolus appears as a hyperintense dot in all three examples, in agreement with prior studies (Khimchenko et al., 2018, Eckermann et al., 2021: PNAS). However, the cytoplasm of the neurons showed different contrast with respect to the surrounding tissue. The signal in the cytoplasm was either isointense (Figure 10A, C), hyperintense (Figure 10E) or hypointense (Figure 10F) in comparison with the surrounding tissue. In one case, the edge of the neuron appeared darker than the surrounding and than the cytoplasm, possible due to local shrinkage, similar to what can be observed in blood vessels (Figure 10A). Further investigations are required to identify which intracellular components give rise to such contrast differences (e.g. organelles or cytoplasm). Nonetheless, this comparison of a few neurons already points towards the rich variety of features that can be observed in SR PhC-µCT images at the cellular level.

**Figure 9.**
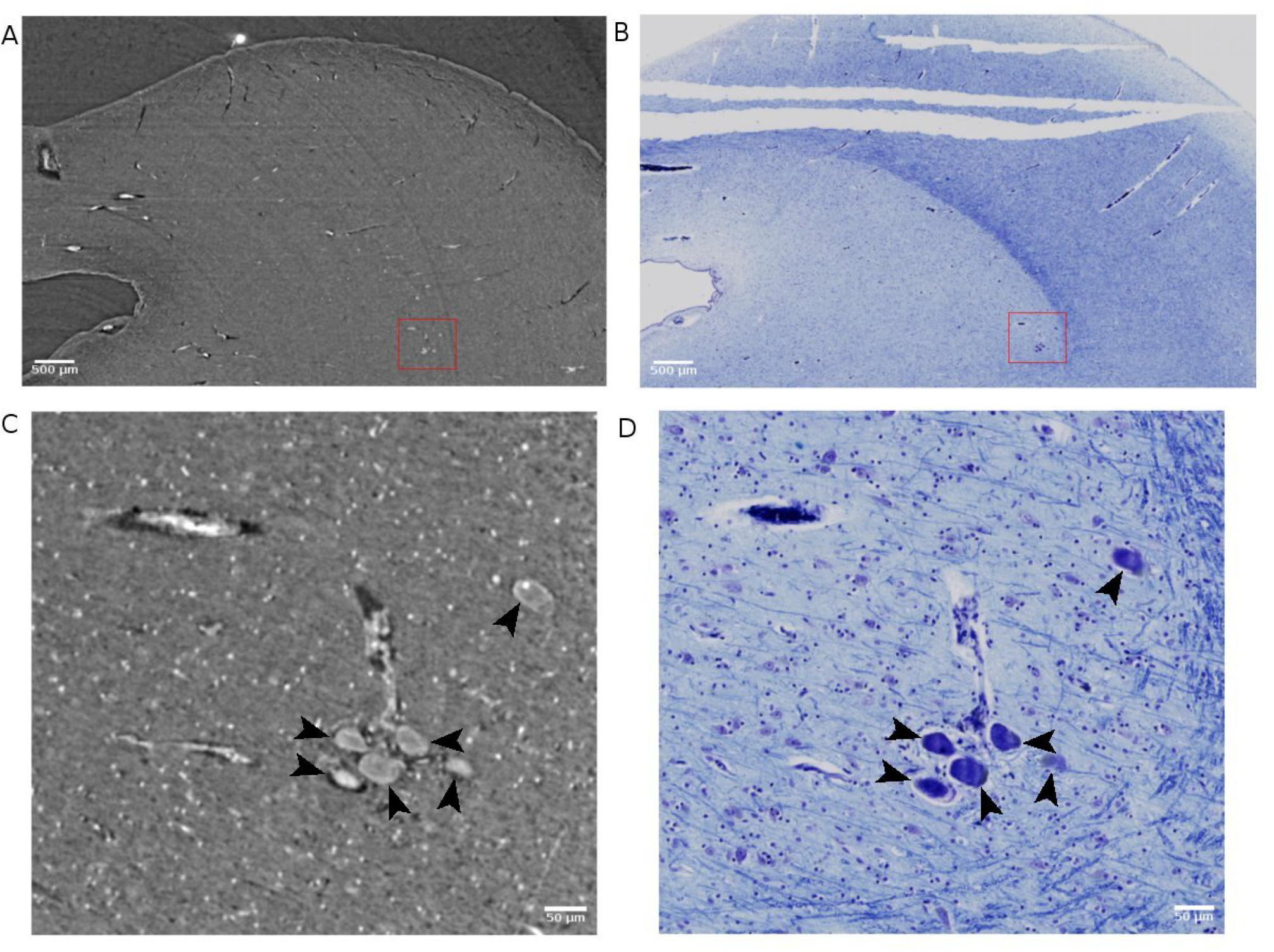
Large neurons detected with synchrotron radiation phase-contrast microtomography (SR PhC-µCT) and histology. (A) SR PhC-µCT acquired with 4.94 µm isotropic voxel size. (B) Histology of a field-of-view (FOV) matching panel A. (C) SR PhC-µCT with 0.94 µm voxel size of the red rectangle region in panel A. Black arrows point to large neurons. (D) Histology of the FOV matching panel C. Scale bars: (A, B) 500 µm, (C, D) 50 µm

**Figure 10.**
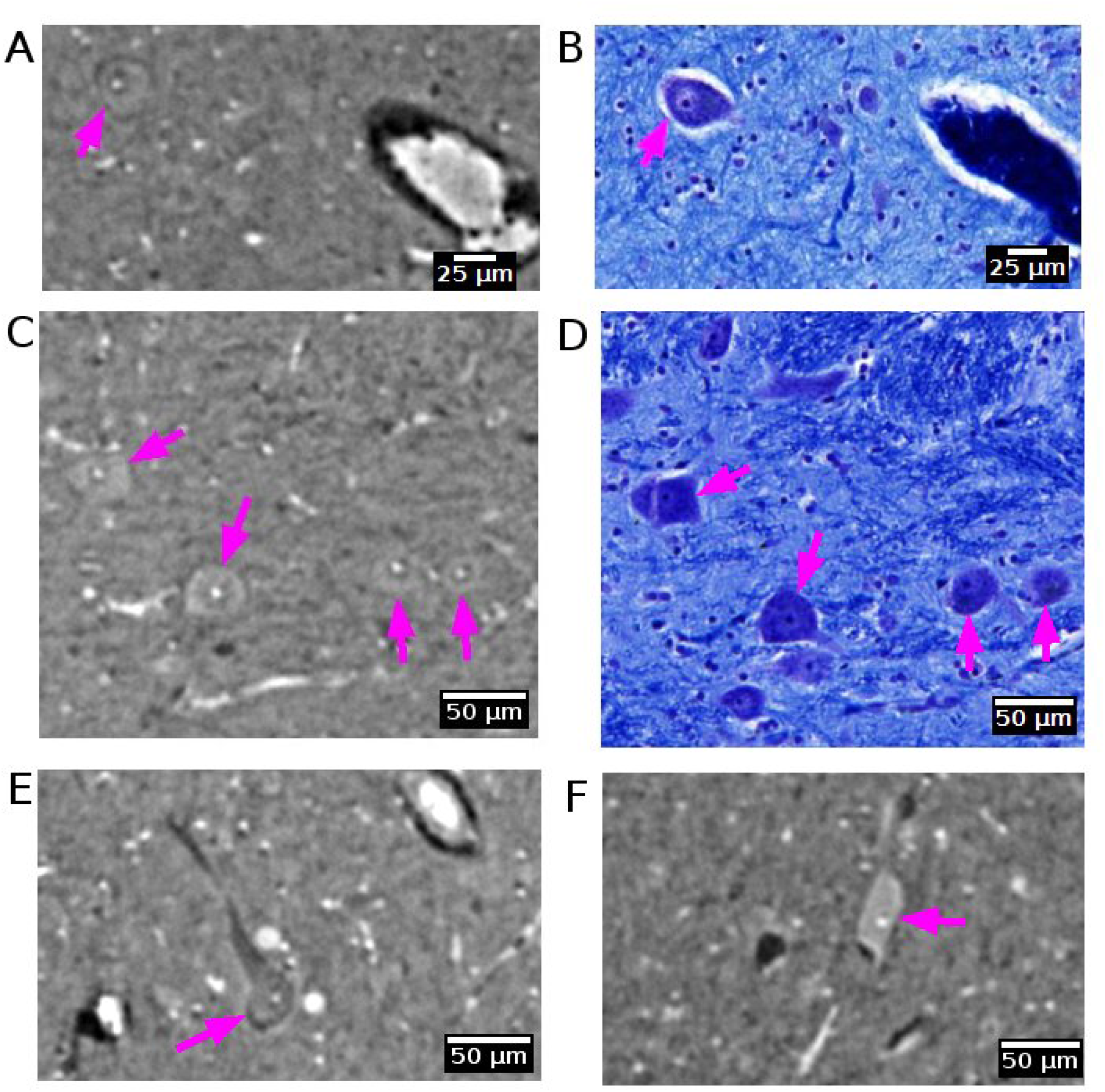
Neurons detected with synchrotron radiation phase-contrast microtomography (SR PhC-µCT). (A-D) Some neurons are found in both SR PhC-µCT and histology. The neuron in panel A shows the same intensity within the cytoplasm as in the surrounding tissue, whereas neurons in panel C show higher intensity in the cytoplasm. (E, F) Examples of neurons with different intensity within the cytoplasm, with lower intensity of cytoplasm shown in panel E and higher intensity observed in the cytoplasm in panel F. Histology of the matching regions were not available. The magenta arrows point to neurons in each panel. SR PhC-µCT are acquired with 0.94 µm isotropic voxel size

## Neuromelanin

Neuromelanin is a dark pigment that progressively accumulate in catecholaminergic neurons, such as dopaminergic neurons in the substantia nigra. In histological slides, neuromelanin appears naturally brown without any staining. It contains several elements with large atomic weight such as iron, zinc and selenium (Bohic et al., 2008). Hence, it is likely that neuromelanin gives rise to higher X-ray attenuation than that typically observed in the cytoplasm of neurons (Figure 11). Indeed, in the substantia nigra we observed very bright spots with shapes that were similar to histology sections of a matching part in substantia nigra. From the single specimen analyzed so far, it appears that the neuromelanin-containing part of the cytoplasm yields the strongest contrast.

**Figure 11.**
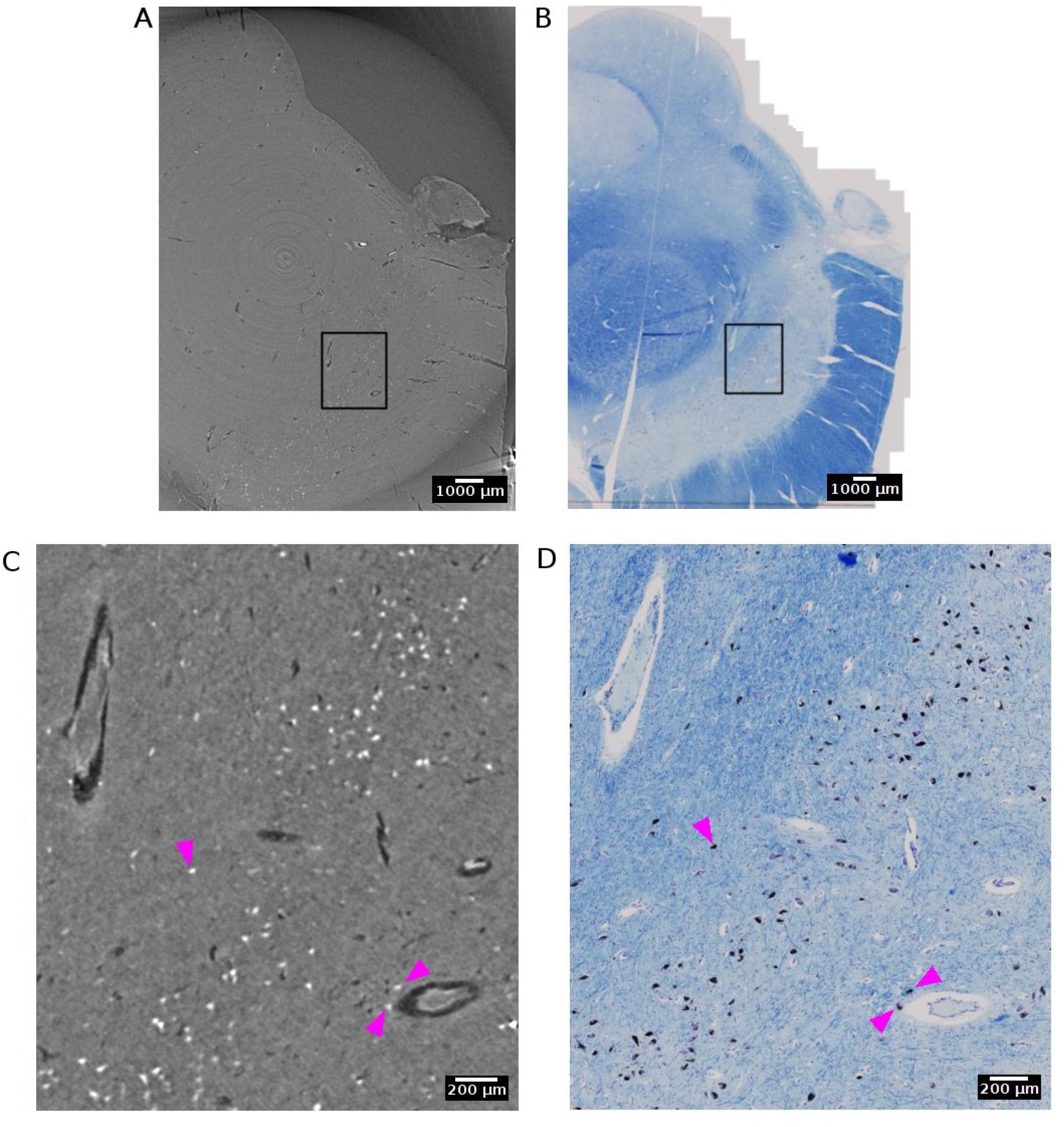
Neuromelanin in the dopaminergic neurons of the substantia nigra detected with synchrotron radiation phase-contrast microtomography (SR PhC-µCT). (A) SR PhC-µCT of the substantia nigra region acquired with 4.74 µm voxel size. (B) The histology of the matching region of panel A. (C) A zoomed inset of rectangle in A. Neuromelanin, annotated with magenta arrows, have outstanding contrast to the surrounding tissue. (D) A zoomed inset of rectangle in B, with neuromelanin annotated with magenta arrows. Scale bars: (A, B) 1 mm. (C, D) 200 µm. through complementary histology, and perhaps preferably electron microscopy, on a small number of co-localized slides.

## Corpora amylacea

Corpora amylacea are polyglucosan granules that are found in multiple organs, including the brain (Riba et al., 2021). They are under active investigation for their relationship to aging, neurodegeneration, immune system and glymphatic transport mechanisms in the brain. The current understanding is that CA is mostly composed of carbohydrates (Sakai et al., 1969). In SR PhC-µCT, these granules can be identified as hyperintense structures with a spherical shape (Fig 12, Fig S4) (Hieber et al., 2016). Recently, a 3D distribution of CA within the human brain stem was analysed from four subjects with mean age of 76 years old, reporting an increased density in the dorsomedial column of the periaqueductal gray (Lee et al., 2023). At the time being, the specificity of our CA detection method must be further investigated, since we anticipate that other densely packed spherical granules encountered in the brain such as Lewy bodies (Koh et al., 2006) may look similar to CA in SR PhC-µCT measurements. In future studies, such uncertainties can be investigated

**Figure 12.**
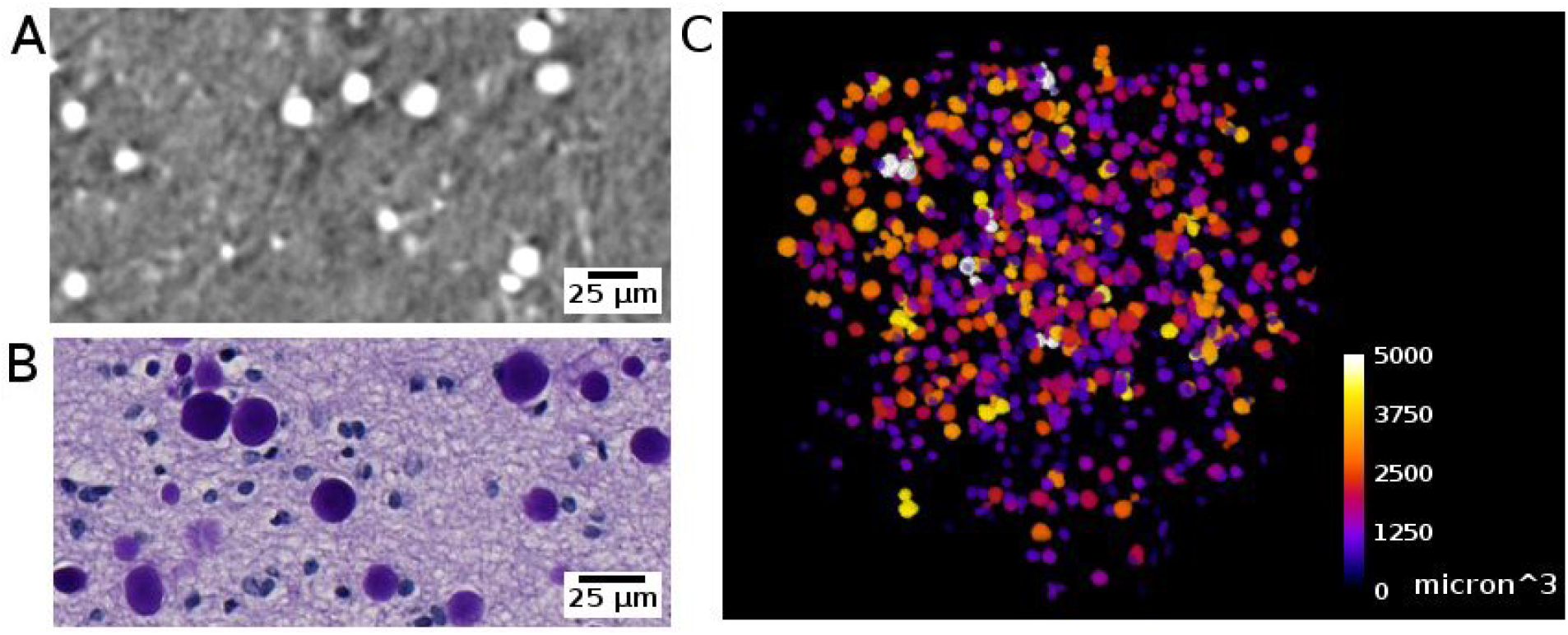
Corpora amylacea detected with synchrotron radiation phase-contrast microtomography (SR PhC-µCT) and histology. (A) Corpora amylace within the human brain tissue is visible in SR PhC-µCT (0.94 µm isotropic voxel). (B) The histology of an adjacent region of panel A. (C) Segmentation result of corpora amylacea from SR PhC-µCT, color-coded by size of the granules.

## Fiber tracts

We were able to observe some of fiber tracts. For example, the oculomotor nerve appeared as hyperintense against the surrounding tissue (Fig 13). Formalin fixed paraffin embedded samples may not be the best option for visualizing fiber structure. In a recent study using mouse brain samples showed that neuronal fibers have higher contrast after ethanol dehydration than after paraffin embedding (Rodgers et al., 2022: *Journal of Neuroscience Methods*). On the other hand, they also observed variations between different fibers. The attenuation coefficient of fibers in cerebellum was lower than fibers in the striatum. Since axonal density, diameter, and myelin thickness can vary considerably between brain regions, we cannot expect fiber tracts to show the same appearance across brain regions. Typically, the myelin thickness increases linearly with increasing axonal diameter. This has been observed both for large white matter fibre tracts in human tissue (Liewald, et al., 2014) and in mouse cortex (Snaidero et al., 2020). Using freeze drying, it has been shown that myelinated axonal tracts in the mouse striatum can be segmented from PhC-μCT images based on their specific attenuation coefficients in this tissue preparation (Mizutani, et al., 2016)

**Figure 13.**
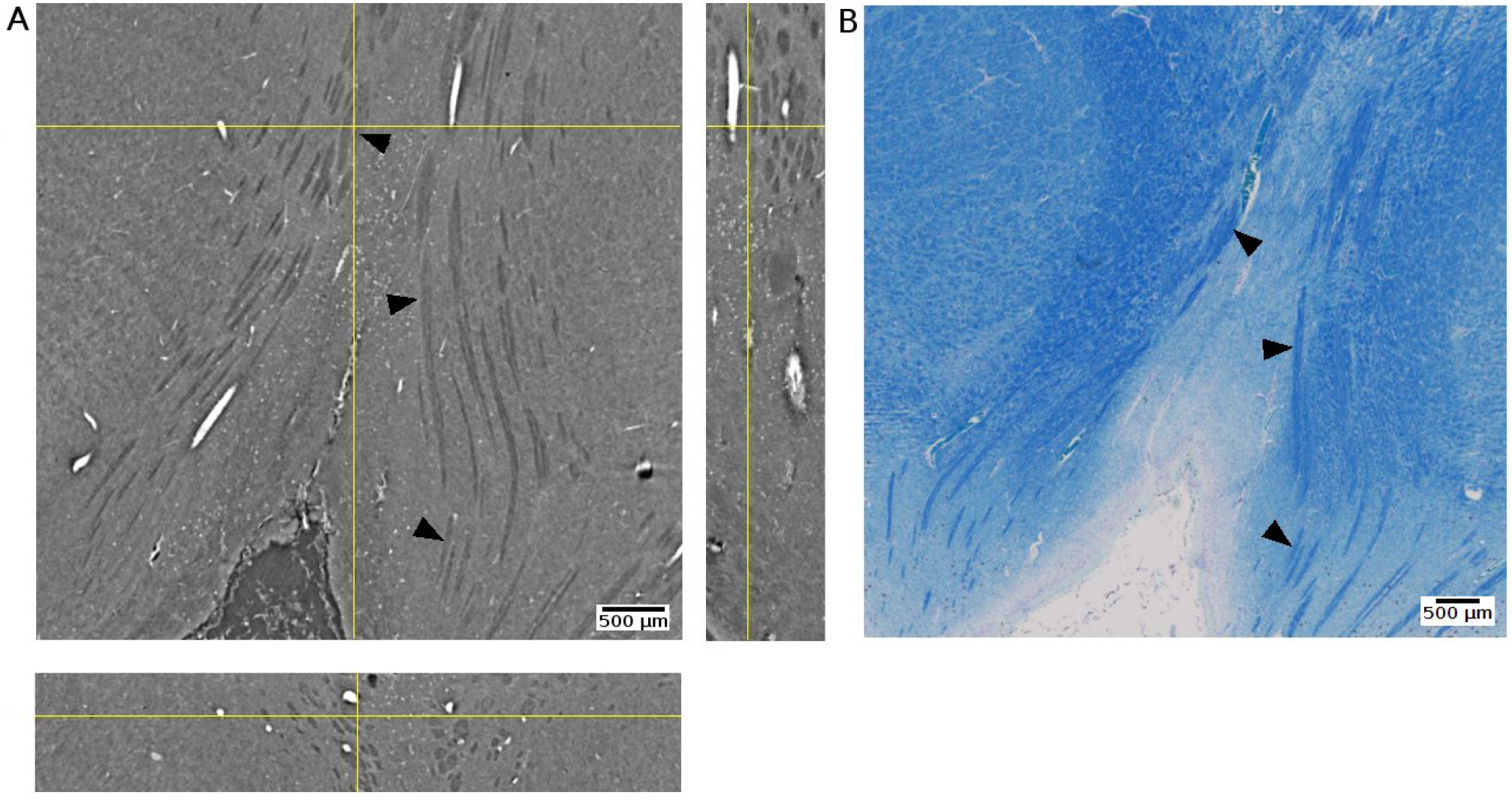
Oculomotor nerve detected with synchrotron radiation phase-contrast microtomography (SR PhC-µCT) and histology. (A) An orthogonal view of the posterior human midbrain acquired with SR PhC-µCT (4.74 µm isotropic voxel). The oculomotor nerve fibers appear as hypointense (B) Histology of the adjacent region stained with luxol fast blue-cresyl violet method. The fiber tracts appear in blue color, since myelin is stained with the luxol fast blue stain. Arrows point to examples of oculomotor nerve fibers in both imaging modalities.

## Discussions

This paper provides a step-by-step guideline for imaging unstained FFPE human brain tissue samples using SR PhC-µCT starting from tissue preparation and measurement to image processing. Although the protocol is written based on the SYRMEP beamline, it can also serve as a general guideline for potential users of other imaging beamlines. Reconstruction and post-processing workflow focus on graphic-user-interface based methods (e.g. STP, ImageJ, ITK-SNAP) to increase accessibility for a broad group of potential users. We also present an overview of biological features that can be identified from the unstained human brain ranging from blood vessel networks to intracellular components such as neuromelanin.

The protocol can be modified to enable particular research goals. Besides the propagation-based method, there are other phase contrast methods, such as grating interferometry or edge illumination techniques, that might be applied for human brain imaging. The grating interferometry provides more reliable estimates of refraction but requires longer measurement times and a more complex reconstruction process (Lang et al., 2014, Astolfo et al., 2016). Edge illumination has potential for soft tissue imaging even in a laboratory setup but also requires longer measurement time (Olivo et al., 2021). These options may be considered for obtaining phase-contrast imaging of soft tissue using conventional X-ray systems. For the segmentation approach, we point towards the use of intensity threshold and morphometry based methods. On the other hand, machine learning segmentation methods such as U-Net (Falk et al., 2019) or the random tree algorithm (Arganda-Carreras et al., 2017) may be useful, if enough training data are available.

With the wide range of new brain imaging methods under development, there is a need for a gold standard method. Researchers within the neuroscience community are beginning to endorse phase contrast X-ray microtomography as a putative novel gold standard for structural imaging (Andersson et al., 2020, Kuan et al., 2020), along with cryo-electron microscopy. For example, microtomography has been used to validate angiography acquired from ultrasound localization microscopy (Chavignon et al., 2021). Furthermore, SR PhC-µCT can show sub-cellular morphologies of neurons with no labeling (Khimchenko et al., 2018).

Synchrotron radiation X-ray is unique for its light with high flux and spatial coherence, enabling the application of phase contrast techniques with fast acquisition times. However, it is limited by the scarce availability of sources, which leads to restricted access. In order for synchrotron radiation light sources to best serve science, users should be encouraged to make data available as open source (Bicarregui et al., 2015). It will reduce possible redundancy and allow acquisition of more samples by sharing of the workload between research groups. Some facilities such as the European Synchrotron Radiation Facility already provide user friendly portal services for accessing open data.

Developments in laboratory based PhC-µCT are projecting new possibilities as well. Microfocus X-ray sources from liquid-metal jet sources provide partially coherent light that enables phase contrast imaging. The feasibility for imaging down to the cellular level has already been demonstrated (Töpperwien et al., 2018: PNAS). If the time required for measurement and reconstruction is reduced to below one hour, the technique may even enable intra-operative imaging (Partridge et al., 2022). Success of surgical removal of tumor cells may be assessed during surgery by imaging of the resection margins (Twengström et al., 2022)

The future of SR PhC-µCT for imaging unstained human brain samples has great potential (Stampfl et al., 2023). To the best of our knowledge, there is only one dataset of a whole human brain available at the moment, which has been acquired with 25 µm isotropic voxel size (Walsh et al., 2021). This voxel size allows for identifying large blood vessels and delineating the boundary between grey and white matter. However, it is too large for investigating the brain at a cellular level. Ideally, whole human brain samples should be scanned at high resolution as done recently with the mouse brain (Rodger et al., 2022: SPIE). Higher resolution images combined with automatic cell classification is likely to become a great resource for understanding the human brain at an advanced level of detail (Eckermann et al., 2021: PNAS). With further improvements, SR PhC-µCT has the potential to provide unprecedented insights into human brain anatomy and neurological diseases.

## Author Contributions

JYL: conceptualization, investigation, data curation, methodology, project administration, writing— original draft, and writing—review and editing, SD:conceptualization, investigation, methodology, software, writing—original draft, and writing—review and editing, AM: resources, validation, and writing—review and editing, UM: resources, validation, GT: conceptualization, resources, investigation, EL: resources, investigation, data curation, LD: investigation, SM: investigation, writing—review and editing, TS: resources, JB: funding acquisition, writing—review and editing, KS: funding acquisition, writing—review and editing, RL:conceptualization, investigation, methodology, writing—original draft, writing—review and editing, GH: conceptualization, investigation, data curation, methodology, writing—original draft, writing—review and editing, project administration, supervision, funding acquisition. All authors contributed to the article and approved the submitted version.

## Acknowledgments

We acknowledge Elettra Sincrotrone Trieste for providing access to its synchrotron radiation facilities and we thank Diego Dreossi for the cone beam imaging of our samples. We thank Johannes Steiner and Tatjana Steiner from the body donor program, Jürgen Papp, Andreas Wagner, and Peter Neckel for organ handling and Henry Evrard, Alina Novitska, Mark Bailey and Lucas Gehlhaar for support with the Zeiss Axioscan Microscope. Finally, we would like to express our gratitude to the body donors.

## Data Availability Statement

The original contributions presented in the study are included in the article/Supplementary material, further inquiries can be directed to the corresponding author.

## Ethics statement Acknowledgments

The studies involving humans were approved by Medical Department of the University of Tübingen. The studies were conducted in accordance with the local legislation and institutional requirements.

The participants provided their written informed consent to participate in this study.

## Reference

1. Adler, Daniel H., et al. “Histology-derived volumetric annotation of the human hippocampal subfields in postmortem MRI.” Neuroimage 84 (2014): 505–523.

2. Albers, Jonas, et al. “Elastic transformation of histological slices allows precise co-registration with microCT data sets for a refined virtual histology approach.” Scientific Reports 11.1 (2021): 10846.

3. Amunts, Katrin, et al. “BigBrain: an ultrahigh-resolution 3D human brain model.” science 340.6139 (2013): 1472–1475.

4. Amunts, Katrin, et al. “Julich-Brain: A 3D probabilistic atlas of the human brain’s cytoarchitecture.” Science 369.6506 (2020): 988–992.

5. Arganda-Carreras, Ignacio, et al. “Trainable Weka Segmentation: a machine learning tool for microscopy pixel classification.” Bioinformatics 33.15 (2017): 2424–2426.

6. Astolfo, A, et al. (2016). Amyloid-β plaque deposition measured using propagation-based X-ray phase contrast CT imaging. Journal of synchrotron radiation, 23(3), 813–819.

7. Barbone, Giacomo E., et al. “High-spatial-resolution three-dimensional imaging of human spinal cord and column anatomy with postmortem X-ray phase-contrast micro-CT.” Radiology 298.1 (2021): 135–146.

8. Berg, Stuart, et al. “Ilastik: interactive machine learning for (bio) image analysis.” Nature methods 16.12 (2019): 1226–1232.

9. Bicarregui, Juan, Brian Matthews, and Frank Schluenzen. “PaNdata: open data infrastructure for photon and neutron sources.” Synchrotron Radiation News 28.2 (2015): 30–35.

10. Birkl, Christoph, et al. “Effects of formalin fixation and temperature on MR relaxation times in the human brain.” NMR in Biomedicine 29.4 (2016): 458–465.

11. Bohic, Sylvain, et al. “Intracellular chemical imaging of the developmental phases of human neuromelanin using synchrotron X-ray microspectroscopy.” Analytical Chemistry 80.24 (2008): 9557–9566.

12. Brombal L. Effectiveness of X-ray phase-contrast tomography: effects of pixel size and magnification on image noise. J Instrum. 2020;15(01):C01005–C01005. doi:10.1088/1748-0221/15/01/C01005

13. Brun, Francesco, et al. “Enhanced and flexible software tools for X-ray computed tomography at the Italian synchrotron radiation facility Elettra.” Fundamenta Informaticae 141.2–3 (2015): 233-243.

14. Brun, Francesco, et al. “SYRMEP Tomo Project: a graphical user interface for customizing CT reconstruction workflows.” Advanced structural and chemical imaging 3.1 (2017): 1–9.

15. Brunet, J., et al. “Preparation of large biological samples for high-resolution, hierarchical, synchrotron phase-contrast tomography with multimodal imaging compatibility.” Nature protocols 18.5 (2023): 1441–1461.

16. Bukreeva, Inna, et al. “Micro-morphology of pineal gland calcification in age-related neurodegenerative diseases.” Med Phys. 50.3 (2022): 1601–1613.

17. De Guzman, A. Elizabeth, et al. “Variations in post-perfusion immersion fixation and storage alter MRI measurements of mouse brain morphometry.” Neuroimage 142 (2016): 687–695.

18. Delogu, P., et al. “Optimization of the equalization procedure for a single-photon counting CdTe detector used for CT.” Journal of Instrumentation 12.11 (2017): C11014.

19. Donato, S., Peña, L. A., Bonazza, D., Formoso, V., Longo, R., Tromba, G., & Brombal, L. (2022). Optimization of pixel size and propagation distance in X-ray phase-contrast virtual histology. Journal of Instrumentation, 17(05), C05021.

20. Dullin, Christian, et al. “Multiscale biomedical imaging at the SYRMEP beamline of Elettra-Closing the gap between preclinical research and patient applications.” Physics Open 6 (2021): 100050.

21. Dusek, Petr, et al. “The choice of embedding media affects image quality, tissue R2*, and susceptibility behaviors in post-mortem brain MR microscopy at 7.0 T.” Magnetic Resonance in Medicine 81.4 (2019): 2688–2701.

22. Eckermann, Marina, et al. “Phase-contrast x-ray tomography of neuronal tissue at laboratory sources with submicron resolution.” Journal of medical imaging 7.1 (2020): 013502–013502.

23. Eckermann, M., Van der Meer, F., Cloetens, P., Ruhwedel, T., Möbius, W., Stadelmann, C., & Salditt, T. (2021). Three-dimensional virtual histology of the cerebral cortex based on phase-contrast X-ray tomography. Biomedical optics express, 12(12), 7582–7598.

24. Eckermann, Marina, et al. “Three-dimensional virtual histology of the human hippocampus based on phase-contrast computed tomography.” Proceedings of the National Academy of Sciences 118.48 (2021): e2113835118.

25. Edlow, Brian L., et al. “7 Tesla MRI of the ex vivo human brain at 100 micron resolution.” Scientific data 6.1 (2019): 244.

26. Einarsson, Emma, et al. “Phase-contrast enhanced synchrotron micro-tomography of human meniscus tissue.” Osteoarthritis and Cartilage 30.9 (2022): 1222–1233.

27. Falk, Thorsten, et al. “U-Net: deep learning for cell counting, detection, and morphometry.” Nature methods 16.1 (2019): 67–70.

28. Flint, Jeremy J., et al. “Visualization of live, mammalian neurons during Kainate-infusion using magnetic resonance microscopy.” Neuroimage 219 (2020): 116997.

29. Frost, Jakob, et al. “3d virtual histology reveals pathological alterations of cerebellar granule cells in multiple sclerosis.” Neuroscience (2023)

30. Haddad, Tariq Sami, et al. “Tutorial: methods for three-dimensional visualization of archival tissue material.” Nature protocols 16.11 (2021): 4945–4962.

31. Handwerker, Jonas, et al. “A CMOS NMR needle for probing brain physiology with high spatial and temporal resolution.” Nature methods 17.1 (2020): 64–67.

32. Hieber, Simone E., et al. “Tomographic brain imaging with nucleolar detail and automatic cell counting.” Scientific reports 6.1 (2016): 32156.

33. Howard, Amy FD, et al. “An open resource combining multi-contrast MRI and microscopy in the macaque brain.” Nature Communications 14.1 (2023): 4320.

34. Junemann, O., et al. “Comparative study of calcification in human choroid plexus, pineal gland, and habenula.” Cell and Tissue Research (2023): 1–9.

35. Koh, Seong-Beom, et al. “Phase contrast radiography of Lewy bodies in Parkinson disease.” Neuroimage 32.2 (2006): 566–569.

36. Khimchenko, Anna, et al. “Hard X-ray nanoholotomography: large-scale, label-free, 3D neuroimaging beyond optical limit.” Advanced Science 5.6 (2018): 1700694.

37. Kyrieleis, A., et al. “Region-of-interest tomography using filtered backprojection: assessing the practical limits.” Journal of microscopy 241.1 (2011): 69–82.

38. Lang, S., et al. “Experimental comparison of grating-and propagation-based hard X-ray phase tomography of soft tissue.” Journal of Applied Physics 116.15 (2014).

39. Lecoq, Paul. “Development of new scintillators for medical applications.” *Nuclear Instruments and Methods in Physics Research Section A: Accelerators, Spectrometers*, Detectors and Associated Equipment 809 (2016): 130–139.

40. Lee, Ju Young, et al. “Microvascular imaging of the unstained human superior colliculus using synchrotron-radiation phase-contrast microtomography.” Scientific Reports 12.1 (2022): 9238.

41. Lee, Ju Young, et al. “Distribution of corpora amylacea in the human midbrain: using synchrotron radiation phase-contrast microtomography, high-field magnetic resonance imaging and histology” Frontiers in Neuroscience (2023)

42. Legland, David, Ignacio Arganda-Carreras, and Philippe Andrey. “MorphoLibJ: integrated library and plugins for mathematical morphology with ImageJ.” Bioinformatics 32.22 (2016): 3532–3534.

43. Liewald, D., Miller, R., Logothetis, N., Wagner, H. J., & Schüz, A. (2014). Distribution of axon diameters in cortical white matter: an electron-microscopic study on three human brains and a macaque. Biological Cybernetics. 10.1007/s00422-014-0626-2

44. Liu, Yuedong, Cunfeng Wei, and Qiong Xu. “Detector shifting and deep learning based ring artifact correction method for low-dose CT.” Medical Physics (2023).

45. Mai, Hongcheng, et al. “Whole-body cellular mapping in mouse using standard IgG antibodies.” Nature Biotechnology (2023): 1–11.

46. Marone, Federica, Beat Münch, and Marco Stampanoni. “Fast reconstruction algorithm dealing with tomography artifacts.” Developments in X-ray Tomography VII. Vol. 7804. SPIE, 2010.

47. Mizutani, R., Saiga, R., Ohtsuka, M., Miura, H., Hoshino, M., Takeuchi, A., & Uesugi, K. (2016). Three-dimensional X-ray visualization of axonal tracts in mouse brain hemisphere. Scientific Reports, 6(1), 35061. 10.1038/srep35061

48. Müller, Bert, et al. “X-ray imaging of human brain tissue down to the molecule level.” International Conference on X-Ray Lasers 2020. Vol. 11886. SPIE, 2021.

49. Münch, Beat, et al. “Stripe and ring artifact removal with combined wavelet—Fourier filtering.” Optics express 17.10 (2009): 8567–8591.

50. Naidich, T. P., Duvernoy, H. M., Delman, B. N., Sorensen, A. G., Kollias, S. S., & Haacke, E. M. (2009). Duvernoy’s Atlas of the Human Brain Stem and Cerebellum. Springer Vienna. 10.1007/978-3-211-73971-6

51. Nazemorroaya, Azadeh, et al. “Developing formalin-based fixative agents for post mortem brain MRI at 9.4 T.” Magnetic Resonance in Medicine 87.5 (2022): 2481–2494.

52. Ohnesorge, B., et al. “Efficient correction for CT image artifacts caused by objects extending outside the scan field of view.” Medical physics 27.1 (2000): 39–46.

53. Olivo, Alessandro. “Edge-illumination x-ray phase-contrast imaging.” Journal of Physics: Condensed Matter 33.36 (2021): 363002.

54. Ollion, Jean, et al. “TANGO: a generic tool for high-throughput 3D image analysis for studying nuclear organization.” Bioinformatics 29.14 (2013): 1840–1841.

55. Orhan, Kaan, Karla de Faria Vasconcelos, and Hugo Gaêta-Araujo. “Artifacts in micro-CT.” Micro-computed Tomography (micro-CT) in Medicine and Engineering (2020): 35–48.

56. Paganin, David, et al. “Simultaneous phase and amplitude extraction from a single defocused image of a homogeneous object.” Journal of microscopy 206.1 (2002): 33–40.

57. Park, Juhyuk, et al. “Integrated platform for multi-scale molecular imaging and phenotyping of the human brain.” bioRxiv (2022): 2022–03.

58. Piai, Anna, et al. “Quantitative characterization of breast tissues with dedicated CT imaging.” Physics in Medicine & Biology 64.15 (2019): 155011.

59. Peterzol, A., et al. “The effects of the imaging system on the validity limits of the ray-optical approach to phase contrast imaging.” Medical physics 32.12 (2005): 3617–3627.

60. Preibisch, Stephan, Stephan Saalfeld, and Pavel Tomancak. “Globally optimal stitching of tiled 3D microscopic image acquisitions.” Bioinformatics 25.11 (2009): 1463–1465.

61. Raven, Carsten. “Numerical removal of ring artifacts in microtomography.” Review of scientific instruments 69.8 (1998): 2978–2980.

62. Riba, Marta, et al. “From corpora amylacea to wasteosomes: History and perspectives.” Ageing Research Reviews 72 (2021): 101484.

63. Rodgers, Griffin, et al. “Virtual histology of an entire mouse brain from formalin fixation to paraffin embedding. Part 1: Data acquisition, anatomical feature segmentation, tracking global volume and density changes.” Journal of Neuroscience Methods 364 (2021): 109354.

64. Rodgers, Griffin, et al. “Mosaic microtomography of a full mouse brain with sub-µm pixel size.” Developments in X-Ray Tomography XIV. Vol. 12242. SPIE, 2022.

65. Rodgers, Griffin, et al. “Virtual histology of an entire mouse brain from formalin fixation to paraffin embedding. Part 2: Volumetric strain fields and local contrast changes.” Journal of Neuroscience Methods 365 (2022): 109385.

66. Saccomano, Mara, et al. “Synchrotron inline phase contrast µCT enables detailed virtual histology of embedded soft-tissue samples with and without staining.” Journal of synchrotron radiation 25.4 (2018): 1153–1161.

67. Saiga, Rino, et al. “Brain capillary structures of schizophrenia cases and controls show a correlation with their neuron structures.” Scientific Reports 11.1 (2021): 11768.

68. Sakai, Masao, et al. “Studies of corpora amylacea: I. Isolation and preliminary characterization by chemical and histochemical techniques.” Archives of neurology 21.5 (1969): 526–544.

69. Schindelin, Johannes, et al. “Fiji: an open-source platform for biological-image analysis.” Nature methods 9.7 (2012): 676–682.

70. Schneider, Caroline A., Wayne S. Rasband, and Kevin W. Eliceiri. “NIH Image to ImageJ: 25 years of image analysis.” Nature methods 9.7 (2012): 671–675.

71. Schulz, Georg, et al. “Three-dimensional strain fields in human brain resulting from formalin fixation.” Journal of neuroscience methods 202.1 (2011): 17–27.

72. Schulz, Georg, et al. “Multimodal imaging of human cerebellum-merging X-ray phase microtomography, magnetic resonance microscopy and histology.” Scientific reports 2.1 (2012): 1–7.

73. Shepherd, Timothy M., et al. “Inner SPACE: 400-micron isotropic resolution MRI of the human brain.” Frontiers in Neuroanatomy 14 (2020): 9.

74. Snaidero, N., Schifferer, M., Mezydlo, A., Zalc, B., Kerschensteiner, M., & Misgeld, T. (2020). Myelin replacement triggered by single-cell demyelination in mouse cortex. Nature Communications, 11(1), 4901. 10.1038/s41467-020-18632-0.

75. Stampfl, Anton PJ, et al. “SYNAPSE: An international roadmap to large brain imaging.” Physics Reports 999 (2023): 1–60.

76. Tendler, Benjamin C., et al. “The Digital Brain Bank, an open access platform for post-mortem imaging datasets.” Elife 11 (2022): e73153.

77. Töpperwien, Mareike, et al. “Three-dimensional mouse brain cytoarchitecture revealed by laboratory-based x-ray phase-contrast tomography.” Scientific reports 7.1 (2017): 42847.

78. Töpperwien, Mareike, et al. “Three-dimensional virtual histology of human cerebellum by X-ray phase-contrast tomography.” Proceedings of the National Academy of Sciences 115.27 (2018): 6940–6945.

79. Töpperwien, Mareike, et al. “Contrast enhancement for visualizing neuronal cytoarchitecture by propagation-based x-ray phase-contrast tomography.” NeuroImage 199 (2019): 70–80.

80. Töpperwien, Mareike, et al. “Correlative x-ray phase-contrast tomography and histology of human brain tissue affected by Alzheimer’s disease.” Neuroimage 210 (2020): 116523.

81. Tuzzi, Elisa, et al. “Ultra-high field MRI in Alzheimer’s disease: effective transverse relaxation rate and quantitative susceptibility mapping of human brain in vivo and ex vivo compared to histology.” Journal of Alzheimer’s Disease 73.4 (2020): 1481–1499.

82. Twengström, William, et al. “Can laboratory x-ray virtual histology provide intraoperative 3D tumor resection margin assessment?.” Journal of Medical Imaging 9.3 (2022): 031503.

83. Van Aarle, Wim, et al. “The ASTRA Toolbox: A platform for advanced algorithm development in electron tomography.” Ultramicroscopy 157 (2015): 35–47.

84. Van Nieuwenhove, Vincent, et al. “Dynamic intensity normalization using eigen flat fields in X-ray imaging.” Optics express 23.21 (2015): 27975–27989.

85. Vo, Nghia T., et al. “Reliable method for calculating the center of rotation in parallel-beam tomography.” Optics express 22.16 (2014): 19078–19086.

86. Vo, Nghia T., Robert C. Atwood, and Michael Drakopoulos. “Radial lens distortion correction with sub-pixel accuracy for X-ray micro-tomography.” Optics express 23.25 (2015): 32859–32868.

87. Walsh, C.L., Tafforeau, P., Wagner, W.L. et al. Imaging intact human organs with local resolution of cellular structures using hierarchical phase-contrast tomography. Nat Methods (2021).

88. Wang, Ge. “X-ray micro-CT with a displaced detector array.” Medical physics 29.7 (2002): 1634-1636.

89. Wälchli, Thomas, et al. “Hierarchical imaging and computational analysis of three-dimensional vascular network architecture in the entire postnatal and adult mouse brain.” Nature protocols 16.10 (2021): 4564–4610.

90. Wehrl, Hans F., et al. “Assessment of murine brain tissue shrinkage caused by different histological fixatives using magnetic resonance and computed tomography imaging.” (2015).

91. Wehrse, Eckhard, et al. “Ultrahigh resolution whole body photon counting computed tomography as a novel versatile tool for translational research from mouse to man.” Zeitschrift für Medizinische Physik 33.2 (2023): 155–167.

92. Woelfle, Sarah, et al. “CLARITY increases sensitivity and specificity of fluorescence immunostaining in long-term archived human brain tissue.” BMC biology 21.1 (2023): 1–28.

93. Xue, Songchao, et al. “Indian-ink perfusion based method for reconstructing continuous vascular networks in whole mouse brain.” PloS one 9.1 (2014): e88067.

94. Yushkevich, Paul A., et al. “User-guided 3D active contour segmentation of anatomical structures: significantly improved efficiency and reliability.” Neuroimage 31.3 (2006): 1116–1128.

95. Zhanmu, Ouyang, et al. “Paraffin-embedding for large volume bio-tissue.” Scientific reports 10.1 (2020): 12639.

