## Supplementary figures for "Protocol for 3D Virtual Histology of Unstained Human Brain Tissue using Synchrotron Radiation Phase-Contrast Microtomography"

**** Equal contribution as last authors**

### 1 Supplementary Figures

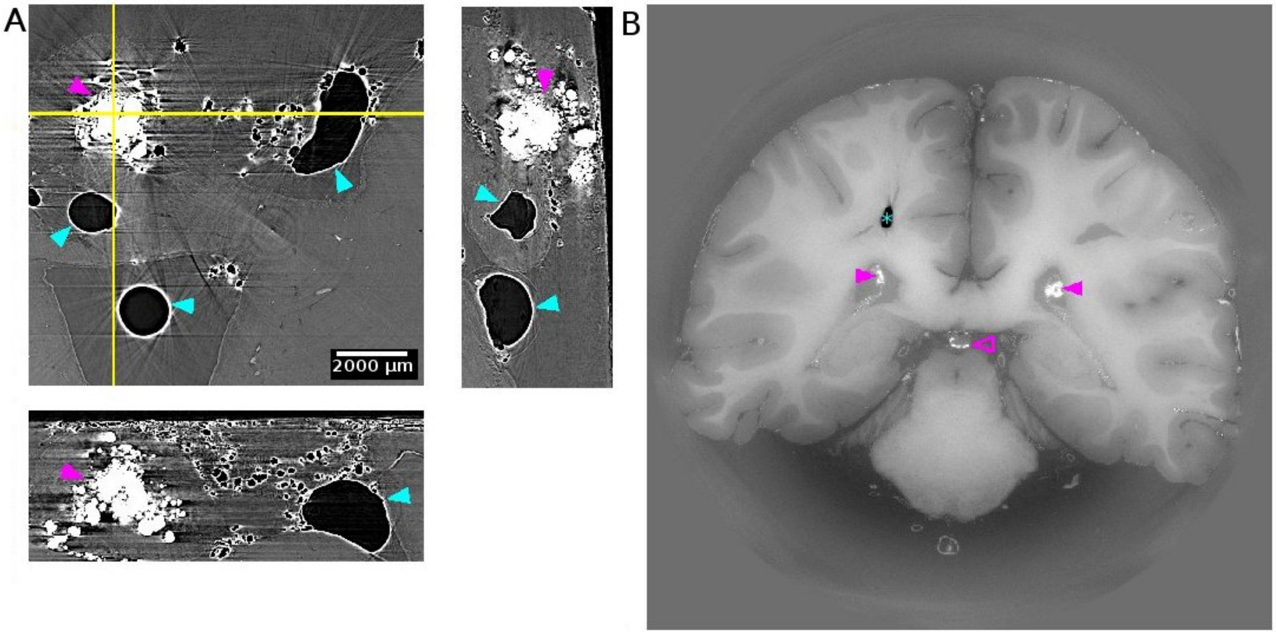


**Supplementary Figure 1. Pineal gland calcification and air bubble.** (A) our data with 4.94 um isotropic voxel size measured at Elettra SYRMEP beamline. Pineal calcification is marked with magenta arrow heads and entrapped air bubbles are marked with cyan arrow heads. (B) Human organ atlas data with 25 um isotropic voxel size measured at ESRF BM05 beamline. Air bubble is marked with asterisk and calcifications are marked with arrows. The filled arrows point to choroid plexus and outlined arrow points to pineal gland. (DOI: 10.15151/ESRF-DC-572252655).

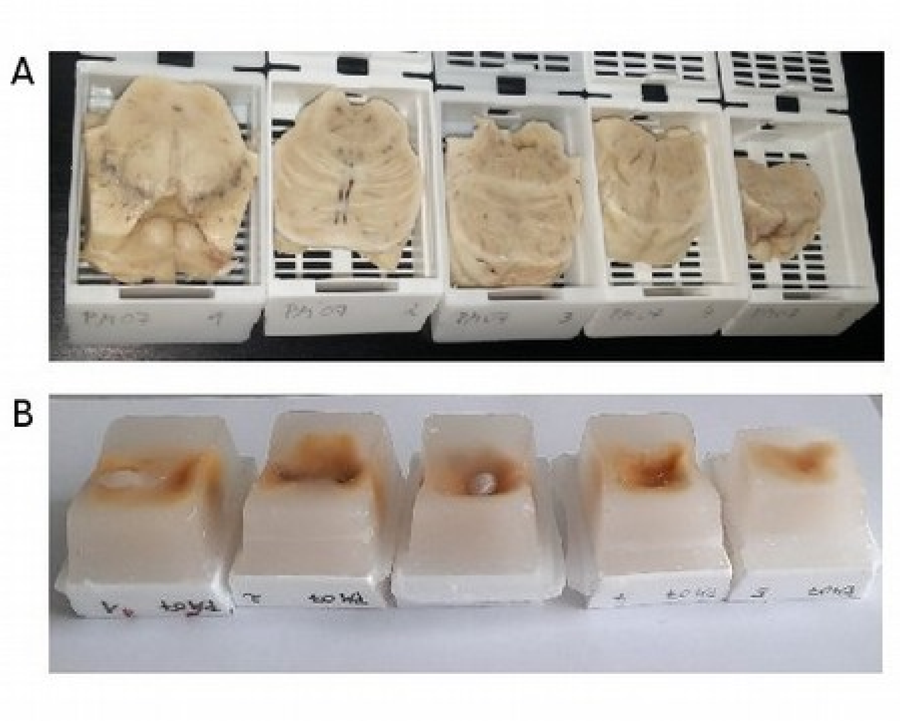


**Supplementary Figure 2. Embedding large samples in paraffin.** (A) Human brain stem samples were cut into pieces prior to embedding. (B) Example of unsuccessful embedding. The surfaces were severely dented. Such dented surface didn't occur when the embedding process were done manually as shown in Figure 1.

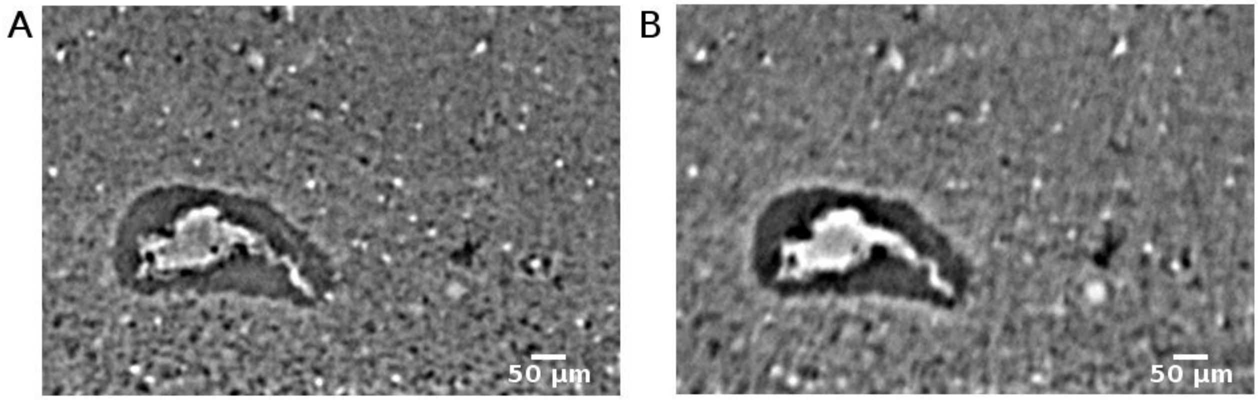


**Supplementary Figure 3. The effect of scintillators thickness.** A tissue was measured with same parameters except the thickness of the scintillator. We used gadolinium gallium garnet Eu-doped scintillators with 17 µm thickness (A) and 45 µm thickness (B). The voxel size is 2 µm istoropic.

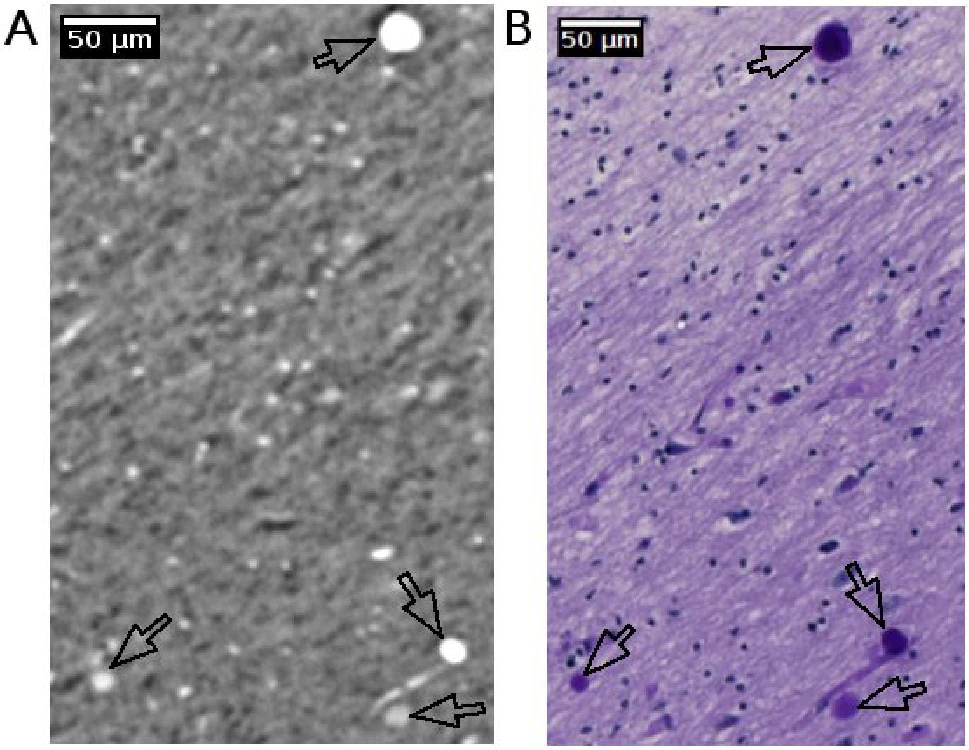


**Supplementary Figure 4. Corpora amylacea in human brain.** Matching regions of the human brain sample with several corpora amylacea imaged with phase-contrast microtomography (A) and periodic acid Schiff stained histology (B). The intensity variance is shown between different granules.

### 2 Supplementary video

**Video 1. Blood vessels in the phase contrast microtomography.** The appearance of the blood vessels in unstained human brain tissue scanned with 0.94 µm istropic voxel.
